# How to characterize a strain? The neglected influence of clonal heterogeneity on the phenotypes of industrial *Saccharomyces*

**DOI:** 10.1101/2020.07.19.211094

**Authors:** Hanna Viktória Rácz, Fezan Mukhtar, Alexandra Imre, Zoltán Rádai, Andreas Károly Gombert, Tamás Rátonyi, János Nagy, István Pócsi, Walter P. Pfliegler

## Abstract

Populations of microbes are constantly evolving heterogeneity that selection acts upon, yet heterogeneity is non-trivial to assess methodologically. The practice of isolating single cell colonies for establishing, transferring, and using a strain results in single-cell bottlenecks with a generally neglected effect on the characteristics of the strain. We used six industrial yeasts to assess the level of heterogeneity in clonal populations, especially in terms of stress tolerance. First, we uncovered the existence of genome structure variants in available sequenced genomes of clonal lineages of thes strains. Subsequent phenotyping of strains and their newly isolated subclones showed that single-cell bottlenecks during isolation can considerably influence the observable phenotype. Next, we decoupled fitness distributions on the level of individual cells from clonal interference by plating single cell colonies. We used the obtained data on colony area for statistical modeling of the heterogeneity in phenotypes. One strain was further used to show how individual subclonal lineages are remarkably different not just in phenotype, but also in the level of heterogeneity. Thereby we call attention to the fact that choosing an initial clonal lineage from an industrial yeast strain may vastly influence downstream performances and observations on geno- and phenotype, and also on heterogeneity.

## Introduction

Yeasts have played an important role in human societies since their ancient domestication. Their biochemical versatility, tolerance to a wide range of stress factors, and the ease of applying traditional and later molecular strain improvement strategies have only increased their roles in many agricultural and industrial fields (Gallone *et al*. 2016; Gonçalves *et al*. 2016; Barbosa *et al*. 2018; Peter *et al*. 2018; Steensels *et al*. 2019). This industrial applicability is most pronounced in the species *Saccharomyces cerevisiae* that has become unsurmountable in the production of leavened bread, alcoholic beverages, bioethanol, and in modern biotechnology, while also being widely utilized in fields like bioprotection, or food and feed supplements (Legras *et al*. 2007; Peter *et al*. 2018). The species is not merely utilized for industrial fermentations, but may be part of the human microbiome or be used as a probiotic (under the taxonomically obsolete name *S. boulardiĩ*), while in some cases, it also has been reported as an opportunistic human pathogen (Peter *et al*. 2018). Importantly, colonizing and infectious isolates are often derived from commercial probiotic or baking strains, or are known to be members of the wine yeast clade (Zhu, Sherlock and Petrov 2016; Pfliegler *et al*. 2017; Peter *et al*. 2018; Imre *et al*. 2019).

*S. cerevisiae* is known to be a genetically diverse species with dozens of globally distributed or endemic phylogenetic clades, many of which show hallmarks of domestication. A number of clades have become adapted to the production of fermented beverages or foods and these are regarded as prime examples of microbe domestication that quite often led to the existence of polyploid and/or aneuploid lineages (Strope *et al*. 2015; Gallone *et al*. 2016; Duan *et al*. 2018; Peter *et al*. 2018; Steensels *et al*. 2019). The most widespread yeast-fermented product worldwide that uses another yeast „species” is lager beer, where fermentation is carried out by domesticated hybrids of *S. cerevisiae* and *S. eubayanus* (known as *S. pastorianus* and *S. carlsbergensis*) (Gallone *et al*. 2019; Langdon *et al*. 2019; Salazar *et al*. 2019).

The key factors in *S. cerevisiae* becoming so ubiquitous in human-made environments are improved fermentation characteristics, including the utilization of various sugars and prodution of aroma components(Steensels *et al*. 2019; Pontes *et al*. 2020), stress tolerance, and elevated adaptability (e.g. Yue *et al*. 2017; Peter *et al*. 2018; Tattini *et al*. 2019). The species is not merely capable of coping with various stress factors found under industrial circumstances, but it also very quickly adapts to changing environments, a trait of utmost importance in the fluctuating environments of various alcoholic fermentations. *Saccharomyces* species are sexual yeasts, able to utilize meiotic recombination to enhance genetic variability to facilitate adaptation (Mortimer 2000; McDonald, Rice and Desai 2016). However, during most industrial processes, yeasts reproduce mitotically. These clonal populations, however, retain their ability to generate novel variants for selection to act upon, in the form of mutations and genome structure variations (GSV). The latter include ploidy changes, aneuploidies/chromosome copy number variations, loss-of-heterozygosity (LOH), gross chromosomal rearrangements (GCR) and mitotic crossing overs (e.g. van den Broek *et al*. 2015; e.g. Zhu, Sherlock and Petrov 2016; Peter *et al*. 2018). These phenomena can alter their industrial performance and may happen very rapidly (Zhang *et al*. 2016; Kadowaki *et al*. 2017; Morard *et al*. 2019; Gorter de Vries *et al*. 2020; Large *et al*. 2020). Along with point mutations, these GSV events result in clonal populations gradually accumulating differences in various traits, leading to clonal heterogeneity, clonal interference (competition among isogenic asexual lineages) and hence the emergence of so-called subclonal lineages, reminiscent of the experimental evolution setups conducted with laboratory strains (e.g. Lang, Botstein and Desai 2011; Payen *et al*. 2016; Blundell *et al*. 2018; Large *et al*. 2020). These evolving and competing subclones ultimately determine the fitness and the performance of the industrial strains during technological applications. Clonal interference may later be alleviated by sexual reproduction, but only if the yeasts survive the technological processes and are able to re-colonize the fermentation environment, as happens in traditional wineries (Mortimer 2000; Magwene 2014). Most modern technological protocols, however, completely remove the applied yeast populations, either immediately or after a limited number of repitchings (Large *et al*. 2020), and new fermentations are carried out with fresh inocula from established propagation companies, e.g. starter cultures in wineries (Ciani *et al*. 2016) or beer yeast starters (Large *et al*. 2020).

In spite of the considerations above, a yeast strain is in general treated as a uniform entity, both in studies aiming at assessing the diversity and characteristics of the species, and in the commercialization and handling of industrial starter yeasts. In fact, strains are by definition genetically uniform microbial cultures. These strains, upon transfer from one lab to another, or even before each experimental round in the same lab, are conventionally spread on agar media to isolate genuine single-cell colonies void of any potential contaminants. Single cell colonies are conventionally considered to be genetically identical (e.g. Eyler 2013) and even in experimental evolution setups, heterogeneity is only considered after the start of the experiment (for a review on ale and lager experimental evolution, see Gibson *et al*. 2020). In the present study, we aimed to investigate whether the wide-spread process of isolating single-cell colonies (subclones) from commercial yeast products and from strains in collections has a hitherto neglected effect on the observable geno- and phenotype of these strains. In particular, as industrial yeasts are propagated en masse (under relatively stressful conditions) by companies producing and packaging them for dozens of generations (Qiu *et al*. 2019; Large *et al*. 2020), we hyphothesized that standing genetic variation and clonal heterogeneity stemming from mutations and genome structure variations may already be present in commercial products and may have considerable effects on the phenotypes of industrial yeast. We also assumed that such a diversity in subclone lineages may confer plasticity to the industrial yeast population as a whole, manifesting in clonal phenotypic heterogeneity. Heterogeneity may presumably cause unpredictable biases to geno- and phenotypic studies involving yeast lineages that need to be isolated from products, whether for basic research, for strain improvement, or for health issues (compare to Pfliegler *et al*. 2017; Large *et al*. 2020).

To observe and compare heterogeneity, we used wine, ale, lager, probiotic, and bread yeasts of various ploidies to study how heterogeneous these yeasts are when colony phenotypes and stress tolerance are considered. Using a baker’s yeast sample, we assessed how even a single population bottleneck, namely the first instance of single-colony isolation, may considerably influence the observed phenotypic characteristics and also the observable clonal heterogeneity of a given strain. Additionally, we discuss that recent genomic studies of yeasts occasionally investigated very different subclone lineages from a single strain, corroborating the widespread nature of heterogeneity in industrial lineages.

## Materials and Methods

### Strains and (sub)culturing

We obtained commercially available ale, bakery, bioethanol, lager, probiotic, and wine yeasts (Table 1.) and pre-cultured the products in YPD medium (VWR, Radnor, PA, USA) for 30 min at room temperature. The bioethanol PE-2 strain was also obtained from NCYC, and the supplier’s protocol was followed for reviving. These cultures were immediately used to isolate 12 subclone lineages from each yeast, which were designated with letters from ’a’ to ’1’.

**Table 1.** Strains used in this study, with accession numbers for whole genome sequencing data.

| Strain name used in this study | Other names | Origin of cultures in this study | Genomes analyzed in this study and their origins |  |  |  |  |
| --- | --- | --- | --- | --- | --- | --- | --- |
|  |  |  | BioSample | SRA Experiment | SRA Run | Coverage (calculated for haploid *S. cerevisiae* genome) | Reference |
| ADY_Baker | Commercial name | Commercial vendor, Hungary, manufactured in Germany, producer undisclosed by vendor | SAMN15579325 (subclone 1) | SRX8770000 | SRR12264806 | 43.0 | This study |
|  |  |  | SAMN15579326 (subclone 2) | SRX8770001 | SRR12264805 | 51.5 | This study |
| Ale | Commercial name | Commercial vendor, Hungary, manufactured in France by a subsidiary of a French company | SAMEA3895632 (isolate sequenced by Peter et al.) | ERX1380630 | ERR1309393 | 30.1 | (Peter *et al.* 2018) |
|  |  |  | SAMN10973883 (isolate sequenced by Langdon et al.) | SRX6781686 | SRR10047300 | 33.8 | (Langdon *et al.* 2019) |
|  |  |  | SAMN10375233 (isolate sequenced by Fay et al.) | SRX4993536 | SRR8173067 | 14.9 | (Fay *et al.* 2019) |
| Bioethanol | PE-2; NCYC 3233, JAY270 (JAY270 is a pure culture isolate derived from a PE2 commercial stock) | NCYC (National Collection of Yeast Cultures) | SAMN04965971 (JAY270 subclone) | SRX2038376 | SRR4047520 | 102.7 | (Rodrigues-Prause *et al.* 2018) |
|  |  | Commercial vendor, Brazil, manufactured in Brazil | SAMN15559291 (product subclone 1) | SRX8748432 | SRR12240130 | 42.3 | This study |
|  |  | Commercial vendor, Brazil, manufactured in Brazil | SAMN15559292 (product subclone 2) | SRX8748433 | SRR12240129 | 30.1 | This study |
|  |  | Commercial vendor, Brazil, manufactured in Brazil | SAMN15559293 (product subclone 3) | SRX8748434 | SRR12240128 | 27.4 | This study |
| Lager | Weihenstephan 34/70 | Commercial vendor, Hungary, manufactured in France by a subsidiary of a French company | SAMN03174146 (isolate A1) | SRX758144 | SRR1649183 | 71.0 | (van den Broek *et al.* 2015) |
|  |  |  | SAMN03174146 (isolate A2) | SRX758149 | SRR1649191 | 87.3 | (van den Broek *et al.* 2015) |
|  |  |  | SAMN03174146 (isolate A1+B11) | SRX758145 | SRR1649190 | 48.2 | (van den Broek *et al.* 2015) |
|  |  |  | SAMD00035489 (isolate sequenced by Okuno et al.) | DRX036594 | DRR040651 | 430.9 | (Okuno *et al.* 2016), also used in (Langdon *et al.* 2019) |
|  |  |  | SAMN10375239 (isolate sequenced by Fay et al.) | SRX4993613 | SRR8172990 | 23.9 | (Fay *et al.* 2019) |
| Probiotic | Commercial name | Commercial vendor, Hungary, manufactured in Hungary with licence from a Canadian company | SAMN11634143 | SRX5874542 | SRR9099591 | 256.8 | (Offei *et al.* 2019) |
| Wine | Commercial name | Commercial vendor (Hungary), manufactured in Switzerland by a subsidiary of a Canadian company | SAMN04286169 | SRX1457336 | SRR2967887 | 20.2 | (Borneman *et al.* 2016) |

### Whole-genome sequencing and ploidy determination

Whole-genome analysis involved previously sequenced genomes downloaded from NCBI SRA. For the Lager and Ale strain, multiple lineages have been sequenced in recent studies, and these were compared (Table 1.). In the case of the ADY_Baker yeast two subclones from a product commercially obtained in Hungary were newly sequenced at the core facility of the University of Debrecen: one typical colony, and one smaller, rough phenotype colony (named subclone 1 and subclone 2, respectively). These lineages were subcultured only once (multiple single-cell bottlenecks were avoided) and were saved as stocks at −70 °C. Genomic DNA was isolated from the lineages after 24 h growth of the cultures following inoculation in the form of a streak on YPD agar from stocks stored at −70 °C. DNA isolation followed Hanna and Xiao (2006). Library preparation was performed using tagmentation with the Nextera DNA Flex Library Prep kit (Illumina, San Diego, CA, USA) according to the manufacturer’s protocol, sequencing was performed using 150 bp paired-end reads on an Illumina NextSeq 500 system, with approximately 50× coverage of the nuclear genome. Altogether three subclones of the Bioethanol strain, obtained from a commercial product in Brazil in active dry yeast form, containing yeast PE-2, were isolated and sequenced at the Bauer Core, Harvard University, Cambridge, MA, using Illumina NextSeq_High 150 paired-end reads. Genomic DNA for these samples was extracted using an in-house protocol, library preparation was carried out using an adapted tagmentation and Nextera kit from Illumina (Baym *et al*. 2015). Raw reads were deposited to NCBI SRA under BioProject PRJNA646688.

Newly generated FASTQ sequencing files along with those obtained from SRA were trimmed and filtered using fastp (Chen *et al*. 2018), and mapped to the S288C reference genome (R64.2.1.) downloaded from the SGD database (yeastgenome.org) and the reference genomes of the other *Saccharomyces* species (Scannell *et al*. 2011) concatenated to it, using bwa 0.7.17. (Li and Durbin 2009). We only used single runs and single experiments for each SRA genome to avoid any effect of clonal heterogeneity in biosamples with multiple available experiments. Sorted BAM files were obtained using samtools 1.7. (Li *et al*. 2009) and Picard-tools 1.124. (http://picard.sourceforge.net) was used to mark duplicated reads. Local realignment around indels and joint variant calling and filtering for the six samples were performed with GATK 4.1.6.0 (Van der Auwera *et al*. 2013; Poplin *et al*. 2018) with regions annotated in the SGD database as simple repeats, centromeric regions, telomeric regions, or LTRs excluded. First, genomic VCF files were obtained, joint calling was applied, and in the resulting VCF files, only SNPs were selected. SNPs were filtered according to the parameters used by (Fay *et al*. 2019): QD < 5.0; QUAL < 30.0; SOR > 3.0; FS > 60.0; MQ < 40.0; MQRankSum < −12.5; ReadPosRankSum < −8.0; --set-filtered-genotype-to-no-call true. Subsequently, biallelic SNPs that were heterozygous in the individual strains were selected and exported to a .csv file using the query option of BCFtools 1.10.2. Here, only those sites were selected that had an allele to allele ratio ≥ 0.2 and at the same time showed an AD value not smaller than one fifth of the strain’s average coverage. Allele frequency plots were obtained from these. Allele frequencies were used to estimate ploidy following Zhu et al. (2016), with the assumptions that diploids have allele ratios of approx. 1:0 or 1:1, triploids of 1:0, 1:2, and 2:1, tetraploids of 1:0, 1:3, 1:1, or 3:1, etc. These ploidy and chromosome copy number variation (CCNV) results obtained from allele ratios were compared to coverage plots and coverage ratios generated by the software Y_MAP_ using chromosome end and GC content bias correction (Abbey *et al*. 2014). In the case of the hybrid W34/70 lager genome, we created a hybrid reference in Y_MAP_ to be able to represent the strain with the same method as well. Results were compared to previous literature on the given strains’ ploidies where available, for Ale (Peter *et al*. 2018; Fay *et al*. 2019; Langdon *et al*. 2019), Bioethanol (Rodrigues-Prause *et al*. 2018), and Lager strains (van den Broek *et al*. 2015; Okuno *et al*. 2016; Fay *et al*. 2019). Called VCF files were uploaded to FigShare (doi: 10.6084/m9.figshare.12673250).

### Multiplex PCR

We performed our recently developed interdelta and microsatellite fingeprinting multiplex PCR method to rule out that the subclones obtained from products are contaminations that do not correspond to the actual strain. Briefly, we combined interdelta, microsatellite (*YLR177w, YOR267c*), and as a control, ITS 1-4 primer pairs in a single PCR reaction (Imre *et al*. 2019). Then, after gel electroforesis we compared the strains to the derived subclones to identify band patterns that could indicate the presence of isolates other than the original strain.

### Colony morphology and petite test

Heterogeneity in colony morphologies (colony phenotype switch) and frequency of *petite* mitochondrial mutants in packed products were assessed by plating samples directly after the first pre-culturing (as described above) onto YPD agar plates (for colony morphologies) and onto GlyYP (glycerol yeast extract peptone) + 0.1% glucose agar plates with cell densities of approx. 200/plate (after cell counting in a haemocytometer). Plates were incubated for 10 days at 30°C (with agar surface facing down) and were visually scored for various phenotypes on YPD (rough, wrinkled, sectored, stalk-like, and very small colonies) and for potential petite mutants on GlyYP. Presumed petites were transferred to YPD and after overnight culturing, were inoculated onto GlyYP plates without glucose. Subclones unable to grow on glucose-free GlyYP were scored as petites. Finally, YPD colonies were washed under tap water to determine the frequency of invasivitiy into agar. At least 1,000 subclone colonies were counted for each strain and for each assay, raw data was uploaded to FigShare (doi: 10.6084/m9.figshare.12673256).

### Spot-plate assays

Tolerance to various stress factors with a focus on industrially relevant stresses (Gibson *et al*. 2007; Qiu *et al*. 2019) and growth on rich and minimal media were assessed using the spot-plate method for all strains and all subclones of the strains. The following stress media based on SD (synthetic defined, 2% glucose, 0.67% yeast nitrogen base without amino acids) were used: ethanol and high sugar osmotic stress (Bioethanol, Ale, Lager, Wine yeasts), NaCl and high sugar osmotic stress (ADY_Baker), and salt and oxidative stress (H_2_O_2_) media for the Probiotic yeast. In preliminary experiments, we determined the optimal concentrations of stressors that may enable differentiating between subclone lineages (Table S1.). Samples grown overnight (30°C) on YPD plates were washed in ddH_2_O, prepared in equal cell concentrations after cell counting with a haemocytometer, and spotted in 10 μl drops in a series of approx. 50,000; 5,000; 500; 50; and 5 cells to the various plates. The samples originating from the initial isolations were briefly stored at 4°C, single-cell bottlenecks were avoided as described above. Plates were incubated at 30°C for 2 days before photographing them using a DSLR camera. Growth was evaluated visually, plate photographs were uploaded to FigShare (doi: 10.6084/m9.figshare.12673253).

### Clonal heterogeneity test

Clonal heterogeneity under stress was assessed by using the same stress conditions as in the spot plate assays, supplemented with assays on SD and YPD media. Freshly grown cells (as described for spot plates) were counted in a haemocytometer and spread to land about 200 cells/plate. Plates were incubated at 30°C for 2 days (YPD and SD media), or for 2, 3, 4, and 6 days (stress media) as colonies reached sizes that were visible but not yet close to each other. For each condition, three replicate plates were used for each sample. Photographs were taken 4 and 6 days after inoculation with a DSLR camera. Data on colony area was gathered by using the Fiji software package CountPHICS (Brzozowska *et al*. 2019) with circularity set to 0.8. Pixel to mm ratios were measured and area calculations were randomly verified by manual measurment in ImageJ for altogether ten colonies (Rueden *et al*. 2017). Plate photographs and all colony area values were uploaded to FigShare (doi: 10.6084/m9.figshare.12673256).

### Statistical analysis of clonal heterogeneity data

Analyses were done in the R environment for statistical computing (R Core Team 2020). Prior to analyzes, colony size data were square root-transformed to bring value distributions closer to Gaussian; also, following square root transformation, data were re-scaled by carrying out z-score transformation (i.e. subtracting variable mean from all values, then dividing by standard deviation) to aid model fitting in later analyzes. We used linear regression modelling of Bayesian approach, utilizing the R-package “MCMCglmm” (Hadfield 2010), because it allows for flexible model specifications, and estimates are less sensitive to group size differences than ordinary least squares methods. Firstly, to test how heterogeneity was dependent on growth conditions across the different strains we fitted a model with the re-scaled colony area measurements as response variable, and specific grouping variable accounting for both strain and growth condition (i.e. practically controlling for strain, condition, and the interaction of these, without including empty factor levels, i.e. untested strain-condition pairs) as fixed predictor. Model specification was done in a way so that group-level residual variances could be estimated. Because measurements originated from Petri dish repeats, and colonies within Petri dishes were of common origin, we included repetition ID nested within strain as random effect to control for non-independence in the data. In the results we assessed growth condition related differences in group heterogeneities by contrasting posterior distributions of residual variance estimates. Statistical significance was established by using 95% highest posterior density (HPD) intervals (analogous to confidence intervals in frequentist modelling): for contrast estimates where the 95% HPD interval did not cross zero, the difference between the contrasted groups is considered to be statistically significant.

Secondly, when testing how subclones differ in colony area heterogeneity from their original sample under salt stress, two separate models were fitted, using data from 4 days and 6 days of incubation. This separate analysis for 4 and 6 days data was necessary because of the non-independence in the data due to the temporal correlation between the measures carried out at day 4 and 6. Since we did not want to test the effect of time (4 *versus* 6 days), and measurements of day 6 inherently depend on (are correlated with) measurements on day 4, using separate models was preferable to more complicated model specifications. We note here that it is possible that a single or multiple subclone lineages within a single colony may appear and quickly invade a sector in a colony, resulting in an asymmetrically growing sectored colony. In such a case, day 6 measurements would not inherently depend on day 4 measurements. However, sectored colonies were found to be very rare even after 10 days of incubation, thus their hypothetical effects can be ignored here. In these models square root- and z-score transformed colony size was the response variable, and strain was fixed predictor. Because we wanted to compare heterogeneities of subclone lineages with that of the commercial ADY_Baker product, in the models residual variances of groups were estimated for strains separately. Similarly to the above described model, repetition ID nested within sample was used as random effect. In the results we report posterior distributions of contrast parameters for residual variance estimates compared between subclones and the initial commercial sample.

For all models weakly informative proper priors were specified; for random effect variances parameter expanded priors were used to aid mixing of the Markov chains for random effect variances. During model fitting, sampling of the posterior distributions were run for 105,000 iterations, from which the first 5,000 were discarded as “burn-in”, and from the Markov chain Monte Carlo (MCMC) process only every 50^th^ samples were retained (called thinning interval), yielding a nominal sample size for parameter estimate posterior distributions equal to 2,000. Model diagnostics included visual checking of MCMC chains for trends in the chain trajectories (plotting MCMC samples in the order of iterations), and calculation of autocorrelation in the MCMC chains at lag of the thinning interval (MCMC chains were considered to be mixing well if absolute value of estimated autocorrelation coefficient was lower than 0.1).

## Results

### Industrial yeast samples and ploidy

In this study, we obtained probiotic, ale, lager, wine, bioethanol, and baking (active dry) yeasts from commercial vendors. Five of these samples belong to strains with sequenced genomes, while the ADY_Baking yeast was sequenced and analyzed in this study for the first time. The genomes of the Probiotic, Bioethanol, and Wine yeast were euploid. The ADY_Baker was euploid tetraploid or aneuploid diploid, depending on the subclone lineage (two of which were sequenced). The Ale and Lager yeast showed previously described extensive aneuploidies, however, these were not identical when different studies were compared and genomes from these were re-analyzed for aneuploidies (Table 2., Figure S1.). Furthermore, strains with multiple sequenced sublineages showed various conspicuous runs of homozygosity (ROH) as well as intrachromosomal changes in coverage pointing to GCR events that often differed between subclones, especially in the case of the Lager yeast (Figure S1.).

**Table 2.** Ploidy and aneuploidies of yeast samples, as found in re-analyzed sequenced genomes and via comparison with literature. Chromosome copy number variations do not include partial (less than half of a chromosome) extra or lost copies arising from segmental duplications or other GCRs which are especially common in the Lager yeasts.

| Strain | Ploidy | Aneuploidy found via genome analysis or described in literature |
| --- | --- | --- |
| ADY Baker | 4 (subclone 1)<br>2 (subclone 2) | no (subclone 1) (This study)<br>yes, also complex rearrangements (subclone 2) +1 I, VI, IX (This study) |
| Ale | 4 (all samples) | yes<br>+1 VI; -1 I, V, XII (Peter *et al.* 2018)<br>-1 I, V (Fay *et al.* 2019)<br>+1 I; +2 VI; -1 XVI (Langdon *et al.* 2019) |
| Bioethanol | 2 (all samples) | no (Rodrigues-Prause *et al.* 2018; This study) |
| Lager | 2+2 hybrid (all samples) | yes, also chromosome segment losses<br>+1 from *S. cerevisiae* II, VII, VIII, X, XI, XIV, XV, hybrid III, *S. eubayanus* VI; +2 from *S. cerevisiae* IX; -1 from *S. cerevisiae* I, VI, *S. eubayanus* IX, X, XV+VIII (A1 van den Broek *et al.*, 2015)<br>+1 from *S. cerevisiae* II, IV, V, VIII, X, XI, XII, XIII, XIV, XV, XVI, *S. eubayanus* X; +2 from hybrid III; +3 from *S. cerevisiae* IX; -1 from *S. cerevisiae* III, *S. eubayanus* I, IX, XV+VIII (A1+B11 of van den Broek *et al.*, 2015)<br>+1 from *S. cerevisiae* II, IV, VII, X, XI, XIII, XIV, XV, hybrid III, *S. eubayanus* VI; +2 from *S. cerevisiae* VIII, IX; -1 from hybrid VII, *S. eubayanus* IX, X (A2 van den Broek *et al.*, 2015)<br>+1 *S. cerevisiae* II, IV, VII, XIII, XIV, XV, XVI, *S. eubayanus* VI; +2 *S. cerevisiae* VIII, IX, hybrid III; -1 *S. eubayanus* II, IX, X (Fay *et al.* 2019)<br>+1 from *S. cerevisiae* I, II, IV, VI, X, XI, XIII, XVI, *S. eubayanus* I; +2 from *S. eubayanus* VI; +3 from *S. cerevisiae* IX, hybrid III; -1 from *S. eubayanus* IV, X, XVI (Okuno *et al.* 2016) |
| Probiotic | 2 | no |
| Wine | 2 | no |

### Heterogeneity of colony phenotypes in commercial yeast products

We determined heterogeneity in colony morphology, invasivity, and *petite* frequency in the industrial yeast strains directly, without subculturing the actual product. We found remarkably variable colony phenotypes (Table 3., Figure 1.) and at the same time, large variability in the fraction of atypical colonies, ranging from 0.78% of variable morphologies (Bioethanol) to as much as 27.36% in the Ale yeasts. In the case of the Ale strain, wrinkled and conspicuously small colonies were the most prevalent. Stalk-like colonies (Figure 1.: m) were observed, although with negligible frequencies, in four of the six strains. Proportion of invasivity among the colonies ranged between 0.31% (Bioethanol) and 34.62% (ADY_Baking), and various types of invasive growth could be observed among the samples. Especially in the case of the ADY_Baking and the Probiotic yeast, different invasive phenotypes co-occured. In the case of the Bioethanol strain, rough morphology and invasiveness always co-occured; in other strains, such a clear link was not observed between these traits. Frequency of *petites* reached more than 1% only in the case of the Ale yeast (Table 3.).

**Table 3.** Frequencies of atypical colony morphologies, invasivity and petite mitochondrial mutants among the strains, tested after minimal pre-culturing of commercial products.

| Strain | Wrinkled curled | Rough undulate | Sectored | Small | Stalk-like | Invasive | *petite* |
| --- | --- | --- | --- | --- | --- | --- | --- |
| ADY_Baking | 0% | 0.72% | 0.72% | 1.22% | 0.14% | 34.62% | 0.00% |
| Ale | 13.98% | 2.13% | 2.96% | 7.98% | 0.30% | 16.57% | 3.44% |
| Bioethanol | 0% | 0.31% | 0.16% | 0.16% | 0.16% | 0.31% | 0.25% |
| Lager | 0% | 7.00% | 2.28% | 0.57% | 0.33% | 0% | 0.41% |
| Probiotic | 0% | 0.45% | 0.68% | 2.15% | 0% | 8.50% | 0.84% |
| Wine | 0% | 0.46% | 0.37% | 1.57% | 0% | 25.97% | 0.47% |

**Figure 1.**
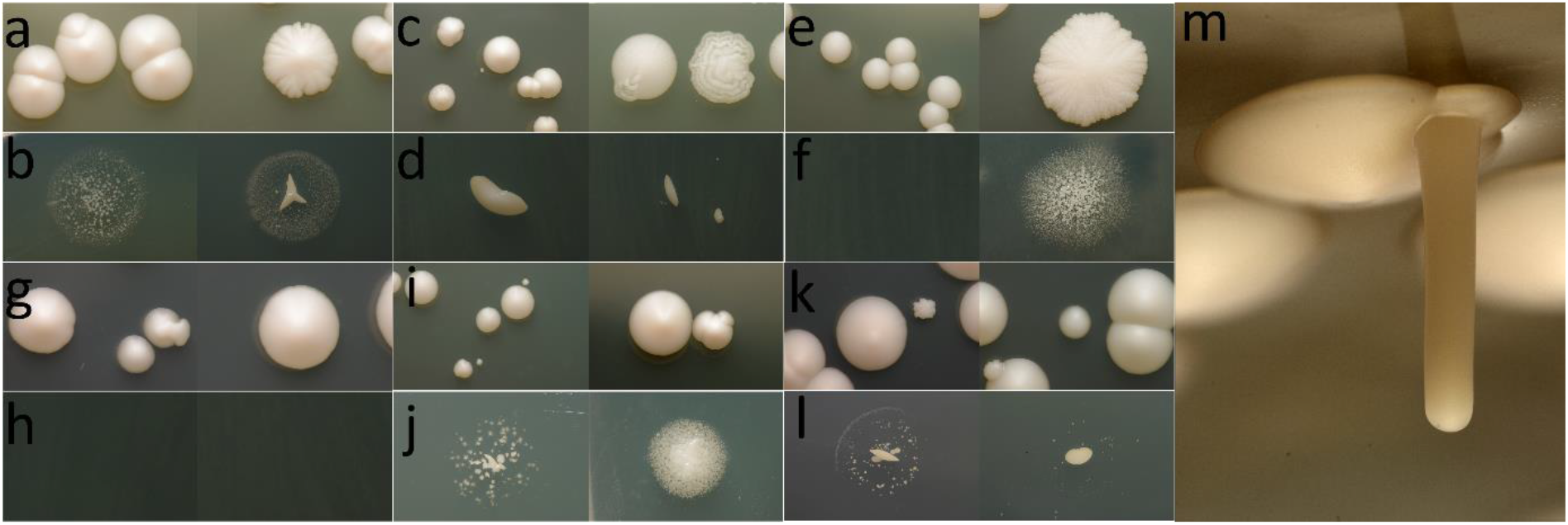
Example colony morphologies (a, c, e, g, i, k, m) and invasivity (or lack of invasive growth) after washing colonies off (b, d, f, h, j, l) observed for industrial yeasts. a-b, m: ADY_Baking, note rough colony on right, and variable invasivity phenotype, along with stalk-like colony; c-d: Ale, note wrinkled undulate and sectored colonies on right; e-f: Bioethanol, note rough invasive colony on right; g-h: Lager, note lack of invasivity; i-j: Probiotic, note very small colonies on left; k-l: Wine, note rough undulate colony on left. Images not to scale.

### Heterogeneity of typical colonies and influence on stress tolerance

After observing heterogeneous colony phenotypes and considerable differences in the frequencies of abnormal colony phenotypes, we isolated 12 subclone colonies from each industrial strain that showed entire, circular, smooth-surface colony phenotypes with the assumption that such regular colonies are the ones most likely to be chosen upon isolation and establishment of a pure lineage in laboratories working with yeasts, while very small or highly unusual colonies are consistently avoided. We avoided subculturing (single-cell bottlenecks) and prolonged culturing of these subclone lineages, and characterized them within weeks of isolation phenotypically, using the colonies saved at 4°C on YPD plates. Thus, we avoided preparing stocks and reviving yeasts from stocks as it has geno- and phenotypic consequences on yeast populations with standing genetic variation (Wing *et al*. 2020). These individual lineages were subjected to multiplex fingerprinting PCR. All subclone lineages showed fingerprinting patterns that were identical, or in the case of the ADY_Baking and Ale yeasts, identical except for the occasional loss of a single band (out of 12 bands). All strains showed clearly different patterns from each other, thus, contamination or cross-contamination of the samples could be excluded and subclone lineages were proven to be derived from the actual strain (Figure S2.).

Spot-plate tests revealed differences in stress tolerance among these subclones lineages established from regular colonies. Visible differences in growth under various stress conditions were observed for half of the strains with the spot-plate method, namely, for the Probiotic, Ale, and ADY_Baker yeast (Table S1.). In all of these cases, a minority of subclone lineages (1–3 subclones depending on strain and condition) showed impaired growth under stress when compared to other subclones or to the original sample that was not subjected to single-cell bottlenecks.

Subsequently, clonal heterogeneity of the six strains during growth on rich and minimal medium and under stress was also evaluated (using colony area as a proxy to fitness). Clonal heterogeneity in the form of variable colony sizes in a single sample from a single strain was prevalent in most samples (Figure 2., S3.). Based on the posterior distributions of residual variance parameters, the strains showed variable levels of heterogeneity in different conditions, which was also apparent from the estimated contrast parameters comparing group-level residual variances between groups (Figure S5.). Group-level residual variances of the measurements estimated with MCMC-GLMM was used to interpret heterogeneity (in this context, higher variation around the group mean corresponds to higher heterogeneity in the measured phenotype, *i.e*. the colony areas). Figure S5. shows that clonal heterogeneity under various growth conditions differs significantly in most cases. That is, when heterogeneities under different conditions were compared, the strains showed significant differences in all (Ale, Lager), or all but one (ADY_Baking, Bioethanol, Probiotic, Wine) of those comparisons. Thus, for every strain, the level of observable heterogeneity was greatly dependent on the condition applied, and the ADY_Baking strain showed the highest differences across conditions. When we compared group heterogeneities (*i.e*. posterior distributions of group residual variances) between strain pairs, separately in each condition (Figure S6.), similarly, the ADY_Baking strain was the yeast that displayed significantly higher measures than others in the highest number of cases, e.g. in minimal medium, its heterogeneity was significantly higher than that of the Ale, Lager, and Bioethanol strain, and statistically not different from that of the Probiotic and Wine yeast. In rich medium, its heterogeneity was significantly higher in all but one pairwise comparison (compared to the Probiotic, its difference was not significant). Under stress conditions (where fewer pairwise comparisons were made due to different stress conditions applied), the Wine and the ADY_Baking strains’ heterogeneities were notable. The former showed significantly lower heterogeneity in three out of four pairwise comparisons, while the latter showed significantly higher heterogeneity in the same number of comparisons.

**Figure 2.**
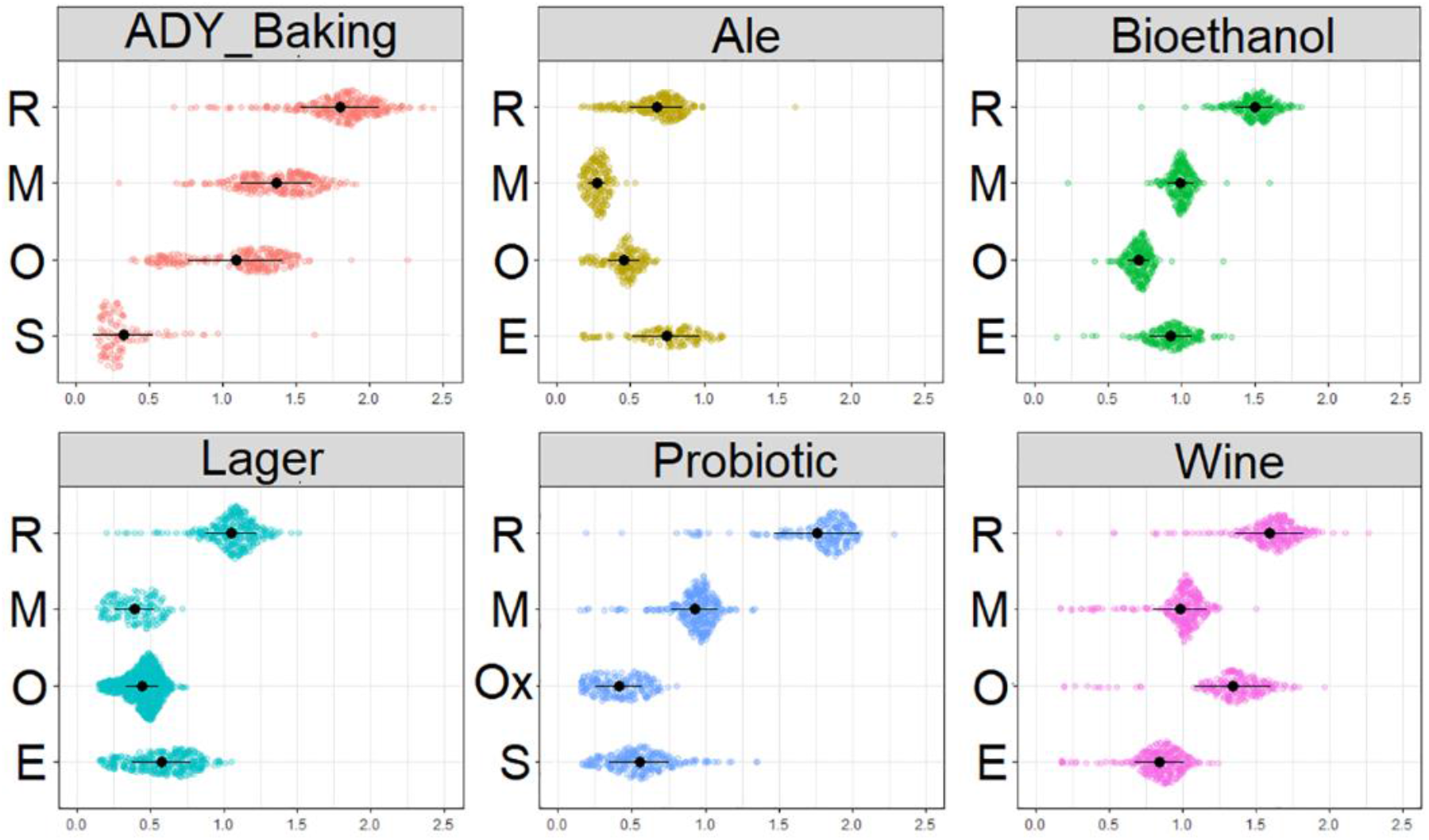
Clonal heterogenity: colony area (square-root transformed) distributions under various conditions for the six commercial samples. R: rich; M: minimal; E: ethanol stress, O: osmotic stress; S: salt stress; Ox: oxidative stress medium. Black dots represent group means, black horizontal lines represent standard deviations.

Based on these results, the ADY_Baking yeast and its subclones were subsequently chosen to further compare how clonal heterogeneity can influence not merely phenotypes but the level of diversity in cell populations derived from subclones. As described above, this strain showed considerable differences in subclones’ spot plate tests (Table S1.), while it showed similarities when two stress conditions were compared (Figure S5.). The level of heterogeneity in the case of subclones and in the original sample under the salt stress condition was compared at two different time points (4 and 6 d) after inoculation, in the following manner. First, growth on minimal SD medium was confirmed to be identical for the subclone lineages using the spot plate method, then the distributions of colony areas were compared under salt stress (Figure 3.). In most cases, heterogeneity was significantly different between the original commercial sample (which generally showed weaker stress tolerance manifesting in generally smaller colonies, but significantly higher heterogeneity) and each of its subclones, except for subclone B (day 4) and subclones B and J (day 6) when mean phenotypes (without residuals) were considered (Figure S7.). Regarding residual variances in colony size distributions, all subclones showed significantly lower heterogeneity compared to the initial commercial sample at both time points except for subclones B, D, and J in the case of day 4 measurements.

**Figure 3.**
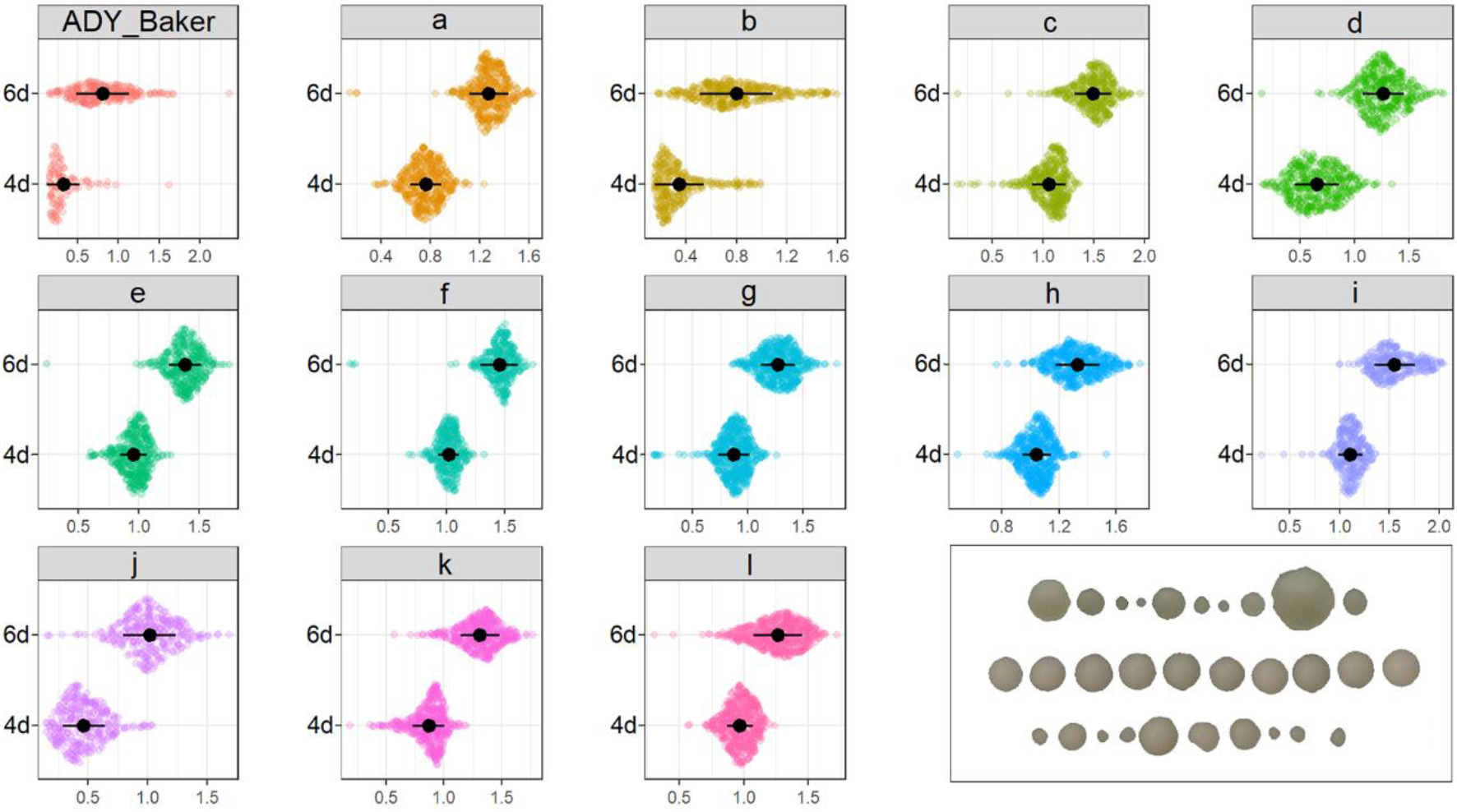
Clonal heterogenity both in growth and in the heterogeneity of growth under stress: colony area (square-root transformed) distributions after 4 and 6 days under salt stress for the ADY_Baking sample and its 12 subclones (named a-l). Black dots represent group means, black horizontal lines represent standard deviations. Inset: illustration of size distributions with ten randomly chosen colonies on the 6th day of incubation for the ADY_Baking yeast (top, note heterogeneity), subclone ’a’ (center, note homogeneity), and subclone ’b’ (bottom, note heterogeneity).

## Discussion

Clonal heterogeneity is a familiar phenomenon for anyone working with culturable microbes. Single-cell isolates from microbial cultures are routinely obtained for various purposes, e.g. for subsequent physiological studies, genetic characterization/modification, metabolic engineering or even for industrial stock propagation, among others, with the advantage of leveraging a simple visual check for eventual contamination with other microbial species. Differences in morphology or size among the grown colonies are often observable to the naked eye. Yet, the underlying causes and, perhaps more importantly, the consequences of single cell bottlenecks (the isolation of a given single-cell colony before an experiment) are mostly neglected.

Studies on the emergence of *de novo* mutations, genome structure variations, and clonal interference in industrial *Saccharomyces* strains (Voordeckers *et al*. 2015; Zhang *et al*. 2016; Bellon *et al*. 2018; Mangado *et al*. 2018; Gorter De Vries *et al*. 2019; Sampaio, Watson and Argueso 2019; Gibson *et al*. 2020; Lairón-Peris *et al*. 2020) have led to increased understanding on their adaptation. In comparison, relatively few yeast studies have been devoted to the importance of clonal heterogeneity in adaptation (e.g. Holland *et al*. 2014; Bódi *et al*. 2017; Vázquez-García *et al*. 2017) or to understanding how epigenetics, gene expression noise, metabolic state, unequal cell division, chronological or replicative age differences, or prions cause yeast populations to be heterogeneous (Halfmann *et al*. 2012; Ackermann 2015; Adamczyk *et al*. 2016; Cerulus *et al*. 2016; Duveau *et al*. 2018). The latter study areas, to our knowledge, exclusively focus on lab strains and not on industrial ones.

Among the factors mentioned above, de novo mutations and GSVs can result in heritable differences among subclone lineages (while other mentioned mechanisms cause constant cell-to-cell heterogeneity without genetic heritability in the strict sense). However, studies comparing *Saccharomyces* strains rarely address the founder effect” of using a subclone lineage of a strain (due to methodological constraints) to characterize the strain itself. Only a few studies have focused on heterogeneous subclone lineages as well as cryptic variation of the PE-2 Bioethanol strain or its derivative JAY270 (Reis *et al*. 2014; Rodrigues-Prause *et al*. 2018; Sampaio, Watson and Argueso 2019) and those of the Lager W34/70 strain (Bolat, Walsh and Turtoi 2008; van den Broek *et al*. 2015), while in most other cases, strains are used interchangeably with subclone lineages. In fact, the commonly used and well-known tetraploid Ale strain of our study, has been sequenced and analyzed by three recent studies, all of which found different karyotypes due to apparent genotypic heterogeneity of the given subclone lineages studied by each (Table 2., Figure S1.). In the case of the tetraploid ADY_Baking active dry yeast, we could identify excessive karyotype heterogeneity within a single batch of the yeast, whic may either be caused by meiotic or mitotic processes. Karyotype changes are important as they are known to be adaptive (Gilchrist and Stelkens 2019) and may even influence cell and colony morphology (Tan *et al*. 2013), and stress adaptations not only in industrial strains (e.g. Kadowaki *et al*. 2017; Morard *et al*. 2019), but in pathogenic *Saccharomyces* as well (Raghavan, Aquadro and Alani 2019).

It must be noted that in the case of pathogenic yeast species, the existence of genotypically different subclone lineages of strains is more often taken into account in the context of comparability among labs (e.g. Franzot *et al*. 1998; Abbey *et al*. 2014) or in the context of heteroresistance to antimycotics (Stone *et al*. 2019). As clinical *Saccharomyces* isolates are regularly derived from commercial (baking and probiotic) yeasts (Pfliegler *et al*. 2017; Imre *et al*. 2019), the clonal heterogeneity inside yeast products should be taken more often into account, when the goal is to understand how stress resistance of industrial yeasts translates into colonizing and pathogenic potential. For example, in a recent study, we compared commercial and clinical yeasts, but did not test multiple subclone lineages of a given strain (Pfliegler *et al*. 2017), a fact that may have influenced our observations due to founder effects.

Based on the facts that genotypic heterogeneity is widespread, especially in tetraploid and hybrid lineages, and that the relatively long (Large *et al*. 2020) industrial yeast cultivations may already be considered a stressful selective environment (Qiu *et al*. 2019), here we designed experiments to quantify and compare heterogeneity in industrial, commercially obtained yeasts. It must be noted that the PE-2 Bioethanol strain was obtained from a culture collection for the phenotyping tets (while its sequenced subclones originated from a commercial product) and thus did not go through extensive culturing before packaging in the form of a yeast product. We found an immense heterogeneity in several cases when colony morphologies and invasivity were assessed inside single batches of one strain (Figure 1., Table 3.). Besides rough, wrinked, and very small colonies, two other observable types are especially interesting. Sectored colonies are themselves naturally arising illustrations of clonal heterogeneity and interference (when lineages inside the colony compete for space as the colony grows), and the fact that in merely 10 days of incubation, sectored colonies were as common as ~2% and ~3% in the Lager and Ale strains, respectively, shows that the emergence of heterogeneous subclone lineages is more of a rule than an exception. The second remarkable colony phenotype was the stalk-like growth previously described and linked to craters in the agar surface by two studies with *Saccharomyces* (Engelberg *et al*. 1998; Scherz, Shinder and Engelberg 2001).

After assessing heterogeneity of single-cell colonies in our strains, we assumed that in routine microbiological workflow, unusual colonies are usually avoided when a pure lineage is to be established. Thus, we obtained 12 subclone lineages that did not show altered morphologies and subsequently showed that even these seemingly uniform lineages can be heterogeneous in their fitness under various stresses (Table S1.). Subsequently, the simple plating method used by us decoupled fitness from clonal interference by isolating cells to form hundreds of distant colonies, enabling the simultaneous study of high- and very low fitness subclone lineages at a given timepoint within a strain or within a subclone lineage. By applying MCMC-GLMM statistic modelling to such single-cell colony measurements, we showed that each strain is different in the level of heterogeneity, while a single strain may also display different levels of heterogeneity depending on the condition (Figure S5-6.). Finally, we also showed that subclone lineages do not only differ in their phenotypes, but can also be significantly different in their potential to generate clonal heterogeneity (Figure S7.). Although we haven’t determined the relative contributions of genetic, epigenetic, or cell age factors affecting heterogeneity, our experimental design of phenotyping (started from overnight cultures on rich media and being evaluated after days of growth on agar media) plausibly strongly suppressed all but the heritable genetic factors. Additionally, the separate growth of colonies on agar media eliminated clonal interference on the test media, enabling the observation of very low fitness lineages emerging from a given strain or a given subclone (Figure 4.).

**Figure 4.**
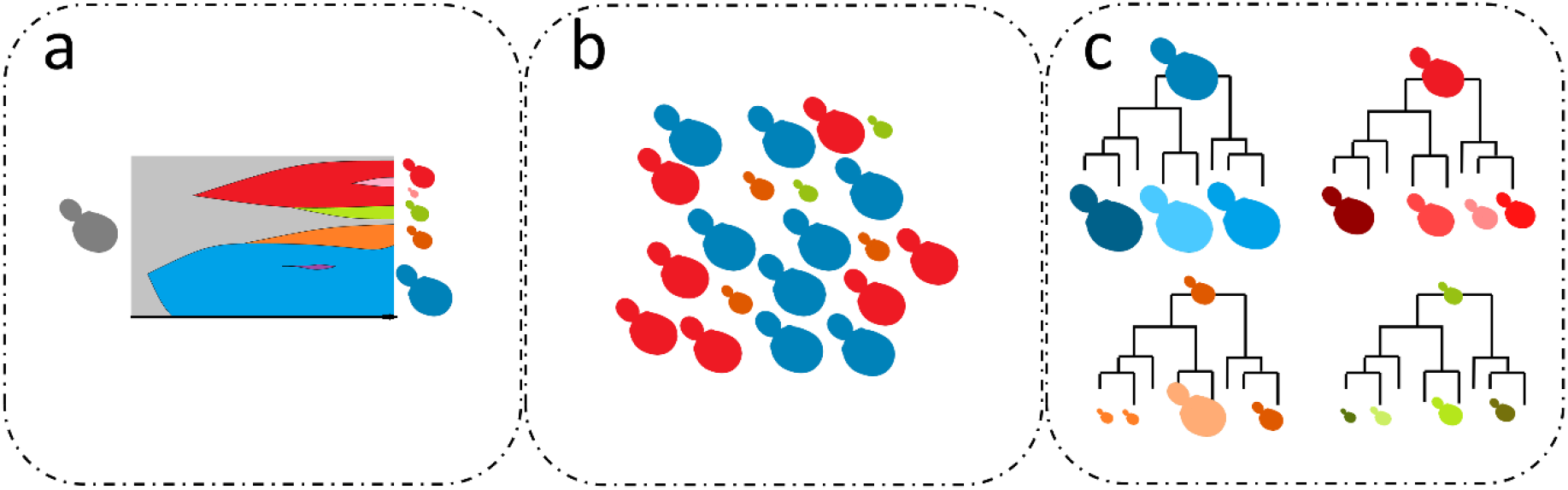
Proposed model of the origin and consequences of clonal heterogeneity in commercial *Saccharomyces* yeasts. a: a stock culture (grey) is used as an inoculum to initiate production, leading to large amounts of cells grown over the course of days (x axis). Subclone lineages (in red, pink, green, orange, and blue) emerge due to genome instability and mutations, compete under (clonal interference) and are selected by stress factors, resulting in changing frequencies of the lineages, as represented on the y axis. b: a final product that contains a heterogeneous yeast population with various frequencies of subclone lineages (clonal heterogeneity). c: subsequent experimental results with single-cell derived subclone lineages lead to a founder effect in the form of different phenotypes and different variability. Colored yeast figure sizes refer to variable fitness, frequency of colors refers to frequency of various lineages in yeast products.

Interestingly, when cells are propagated to be used in an industrial process, these initial propagation conditions can be rather different from the conditions under which the process per se is carried out. Whereas the initial propagation steps have the aim of increasing the microbial population, the process has the aim of generating as much of the product as possible (best TRY compromise; T = Titer, R = rate, Y = yield). Thus, the selective pressure during the propagation step might not only be rather different, but even somehow unfavorable, from the selective pressure during the process itself, i.e. propagation might select subclones that are may not be the best ones for the process.

In conclusion, our experimental setup, to the best of our knowledge, is the first that shows that: 1) clonal heterogeneity is widespread in various clades of commercial yeasts as a presumed consequence of microevolution during the stressful conditions of industrial cell propagation; 2) this heterogeneity affects observable colony morphologies, invasivity, and stress tolerance; and 3) heterogeneity in subsequent generations of a yeast culture is also greatly dependent on which subclone an experiment is based on (summarized in Figure 4.). The surprisingly complex heterogeneity of industrial strains should be taken into account in pheno- and genotyping studies, as well as in strain improvement strategies.

## Acknowledgements

The research is supported by the project “Establishing a scale-independent complex precision consultancy system (GINOP-2.2.1-15-2016-00001)”. We acknowledge support from the Higher Education Institutional Excellence Program (NKFIH-1150-6/2019) of the Ministry of Innovation and Technology in Hungary, within the framework of the Biotechnology thematic program of the University of Debrecen. A.I. was supported by the ÚNKP-19-3-I-234 New National Excellence Program of the Ministry of Human Capacities of Hungary. W.P.P. was supported by a Fulbright Research Award from the Hungarian-American Fulbright Commission. We are deeply grateful for Michael M. Desai, Harvard University, for enabling the genomic sequencing of the three PE-2 clones and agreeing with sharing the corresponding data.

