## Supplementary material for "How to characterize a strain? The neglected influence of clonal heterogeneity on the phenotypes of industrial *Saccharomyces*"

**Supplementary tables and figures.**

**Table S1.** Evaluation of spot plate tests with strains and subclones on rich, minimal, and stress media. ++ strong growth, + medium growth, +- weak growth. Raw images captured for the spot plate tests are uploaded to FigShare.

**Figure S1.** Allele frequency plots (top for each genome, only heterozygous SNPs considered) and chromosome coverage plots obtained with YMAP (bottom for each genome) for the various available sequenced lineages for each strain: ADY_Baking, subclone 1 (this study); ADY_Baking, subclone 2 (this study); Ale (Fay *et al.* 2019); Ale (Langdon *et al.* 2019); Ale (Peter *et al.* 2018); Bioethanol (Rodrigues-Prause *et al.* 2018); Bioethanol subclone 1 (this study); Bioethanol subclone 2 (this study); Bioethanol subclone 3 (this study); j: Lager A1 (van den Broek *et al.* 2015); k: Lager A1+B11 (van den Broek *et al.* 2015), l: Lager A2 (van den Broek *et al.* 2015); Lager (Fay *et al.* 2019); i: Lager (Okuno *et al.* 2016); Probiotic (Offei *et al.* 2019); Wine (Borneman *et al.* 2016). Black dots represent the rDNA tandem repeats (these were masked in allele calling). Note that for Lager yeasts, a composite reference genome was used for mapping, consisting of the two parental species’ reference genomes, although it has been shown that the lager genomes involve major chromosomal rearrangements. Also note that the Probiotic genome sequence has irregular coverage likely resulting from library preparation anomalies. Filtered VCF files used for the allele frequency plots are uploaded to FigShare.

**Figure S2.** Results of multiplex PCR (Imre *et al.* 2019) for the subclones of the six yeast samples used in this study. M: 1 kb DNA size marker.

**Figure S3.** Colony area distributions of commercial strains in different growth conditions. The plot shows the colony area distribution in mm^2^ of the industrial strains examined in different growth conditions. Different strains and media used are represented with colors and letters. Red (a-d): Probiotic strain (YPD, SD, 1mM H_2_O_2_, 2% NaCl, respsectively); light brown (e-h): Bioethanol strain (YPD, SD, 40% glucose, 6% ethanol); dark brown (i-l): Ale strain (YPD, SD, 30% glucose, 4% ethanol); green (m-p): Wine strain (YPD, SD, 30% glucose, 6% ethanol); yellow (q-t): Lager strain (YPD, SD, 30% glucose, 4% ethanol); orange (u-x): ADY_Baking strain (YPD, SD, 30% glucose, 2% NaCl). Raw images used for measurements are uploaded to FigShare.

**Figure S4.** Colony area distributions of the baker strain and its subclones. This plot shows the colony area distribution of the ADY_Baking yeast and its subclones (a-l) after being cultured on 2% NaCl salt stress medium. Measurements were taken after 4 and 6 days (indicated by the colors red and blue, respectively).

**Figure S5.** Graphs showing contrast estimates for pairwise comparisons of various conditions for each of the six strains. Dots show the average difference in residual variance between growth conditions, separately for strains. Horizontal segments show the 95% HPD interval (see Methods); HPD intervals crossing the vertical dashed lines at zero show that in the given comparison the difference is not significant (*i.e.* dots and segments far from zero represent significant group differences). For example, in the ‘ADY_Baking’ strain colonies growing under osmotic and salt stress, variance in the measured phenotype does not differ significantly. However, ADY_Baking yeast colonies on rich medium significantly differ from those grown under salt stress. Note that the contrast parameter in this example is >0, meaning that residual variance (*i.e.* heterogeneity) is higher under the rich condition than under salt stress. R: rich; M: minimal; E: ethanol stress, O: osmotic stress; S: salt stress; Ox: oxidative stress medium.

**Figure S6.** Graphs showing contrast estimates for pairwise comparisons of the six strains under each different condition applied, similarly to Figure S5. Stress 1: osmotic stress. Stress 2: ethanol stress for all pairwise comparisons except for ADY_Baking and Probiotic, where stress 2 refers to salt stress.

**Figure S7.** Graphs showing contrast estimates for pairwise comparisons of the ADY_Baking sample with all of its 12 subclones (a-l) for salt stress condition measured after 4 and after 6 days of incubation. Values higher than 0 correspond to higher variance in the original sample than in the given subclone. Mean phenotypes and residual variances were treated and plotted separately.

**Table S1.**

| Sample | Osmotic (30% glucose) | Salt (2% NaCl) |
| --- | --- | --- |
| ADY_Baker | ++ | ++ |
| ADY_Baker a | ++ | ++ |
| ADY_Baker b | ++ | + |
| ADY_Baker c | ++ | ++ |
| ADY_Baker d | ++ | ++ |
| ADY_Baker e | ++ | ++ |
| ADY_Baker f | ++ | ++ |
| ADY_Baker g | ++ | ++ |
| ADY_Baker h | ++ | ++ |
| ADY_Baker i | ++ | ++ |
| ADY_Baker j | ++ | + |
| ADY_Baker k | ++ | ++ |
| ADY_Baker l | ++ | ++ |
| Sample | Osmotic (30% glucose) | Ethanol (6% ethanol) |
| Ale | ++ | + |
| Ale a | ++ | + |
| Ale b | ++ | + |
| Ale c | ++ | + |
| Ale d | ++ | + |
| Ale e | ++ | +- |
| Ale f | ++ | +- |
| Ale g | ++ | +- |
| Ale h | ++ | + |
| Ale i | ++ | + |
| Ale j | + | + |
| Ale k | ++ | + |
| Ale l | ++ | ++ |
| Sample | Osmotic (40% glucose) | Ethanol (12% ethanol) |
| Bioethanol | ++ | ++ |
| Bioethanol a | ++ | ++ |
| Bioethanol b | ++ | ++ |
| Bioethanol c | ++ | ++ |
| Bioethanol d | ++ | ++ |
| Bioethanol e | ++ | ++ |
| Bioethanol f | ++ | ++ |
| Bioethanol g | ++ | ++ |
| Bioethanol h | ++ | ++ |
| Bioethanol i | ++ | ++ |
| Bioethanol j | ++ | ++ |
| Bioethanol k | ++ | ++ |
| Bioethanol l | ++ | ++ |
| Sample | Osmotic (30% glucose) | Ethanol (6% ethanol) |
| Lager | ++ | ++ |
| Lager a | ++ | ++ |
| Lager b | ++ | ++ |
| Lager c | ++ | ++ |
| Lager d | ++ | ++ |
| Lager e | ++ | ++ |
| Lager f | ++ | ++ |
| Lager g | ++ | ++ |
| Lager h | ++ | ++ |
| Lager i | ++ | ++ |
| Lager j | ++ | ++ |
| Lager k | ++ | ++ |
| Lager l | ++ | ++ |
| Sample | Oxidative (2 mM H_2_O_2_) | Salt (2% NaCl) |
| Probiotic | ++ | + |
| Probiotic a | ++ | + |
| Probiotic b | ++ | + |
| Probiotic c | ++ | + |
| Probiotic d | ++ | + |
| Probiotic e | ++ | + |
| Probiotic f | ++ | + |
| Probiotic g | ++ | + |
| Probiotic h | ++ | + |
| Probiotic i | ++ | + |
| Probiotic j | + | + |
| Probiotic k | ++ | + |
| Probiotic l | + | +- |
| Sample | Osmotic (40% glucose) | Ethanol (12% ethanol) |
| Wine | ++ | ++ |
| Wine a | ++ | ++ |
| Wine b | ++ | ++ |
| Wine c | ++ | ++ |
| Wine d | ++ | ++ |
| Wine e | ++ | ++ |
| Wine f | ++ | ++ |
| Wine g | ++ | ++ |
| Wine h | ++ | ++ |
| Wine i | ++ | ++ |
| Wine j | ++ | ++ |
| Wine k | ++ | ++ |
| Wine l | ++ | ++ |

**Figure S1.**

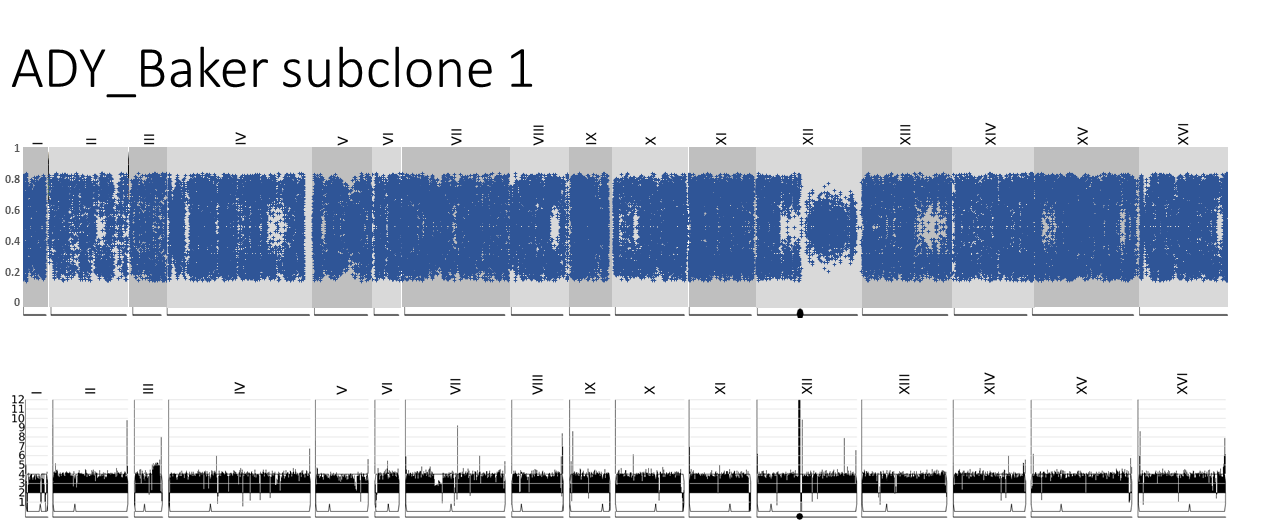

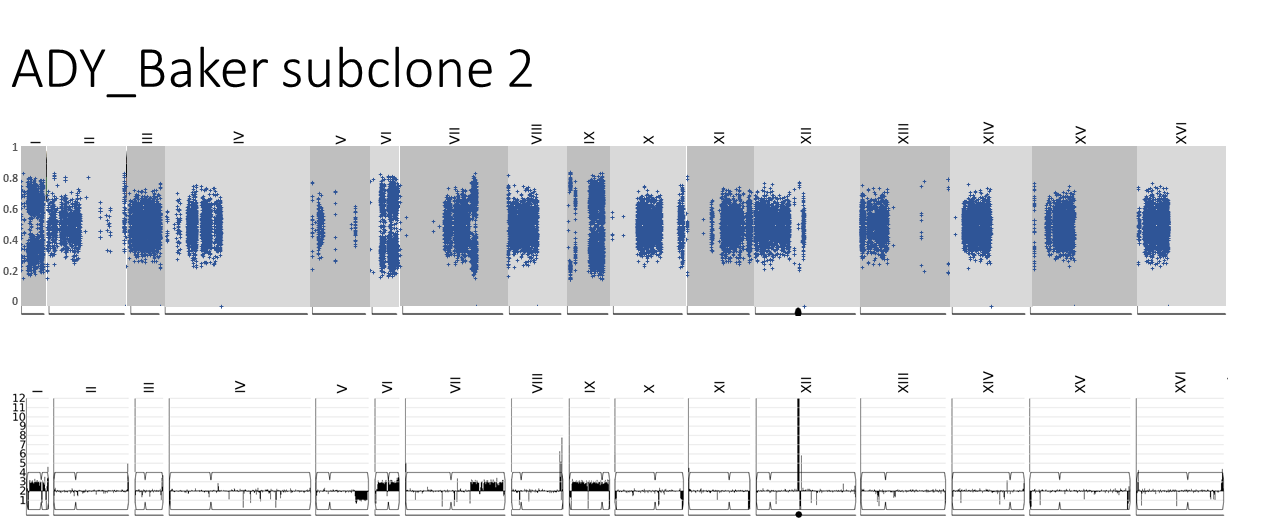

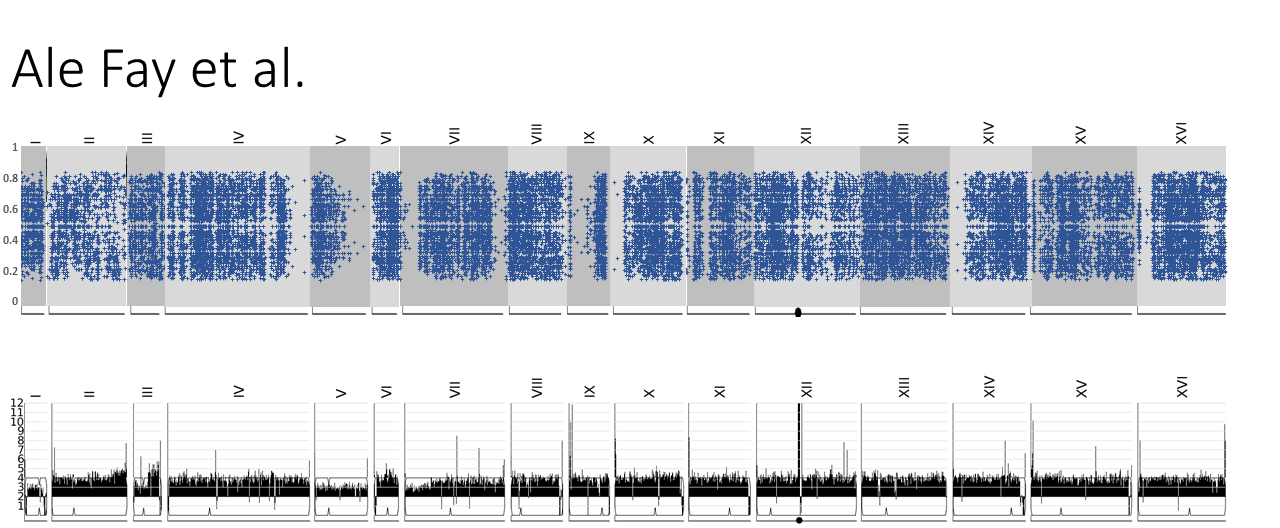

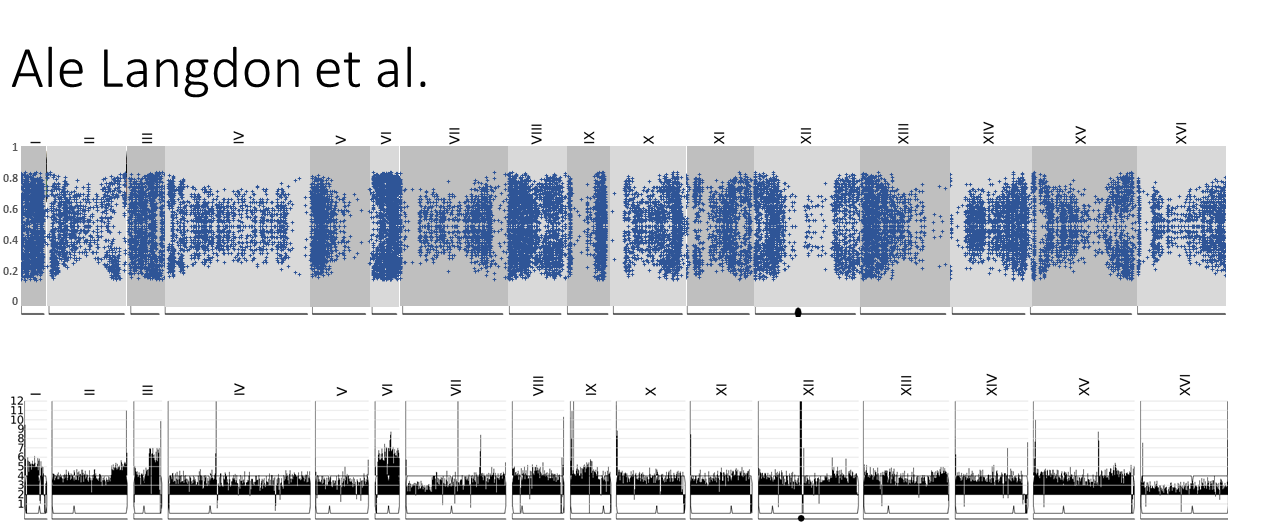

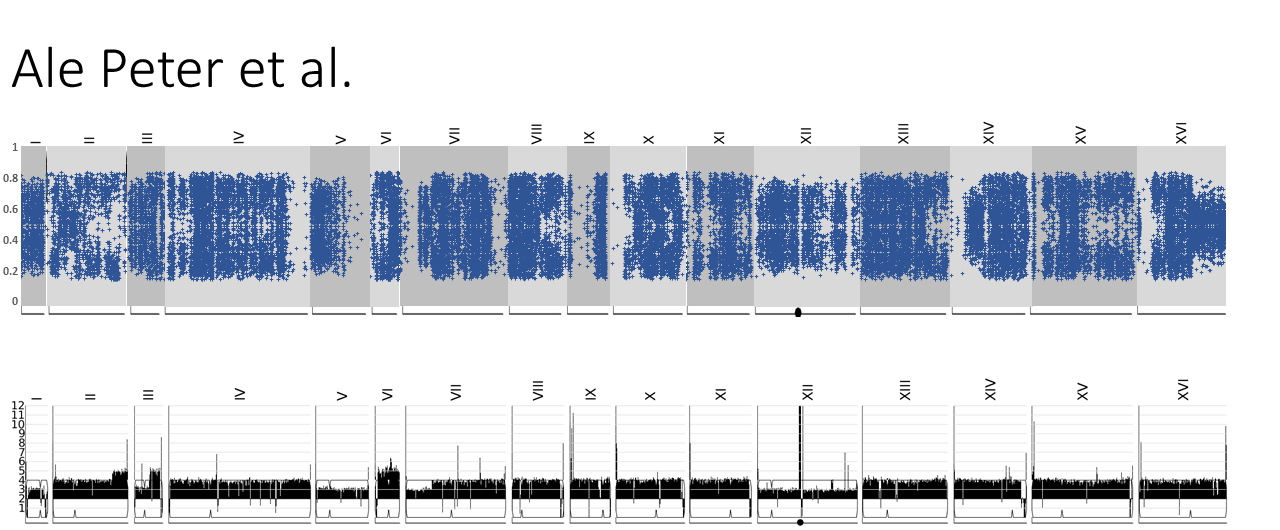

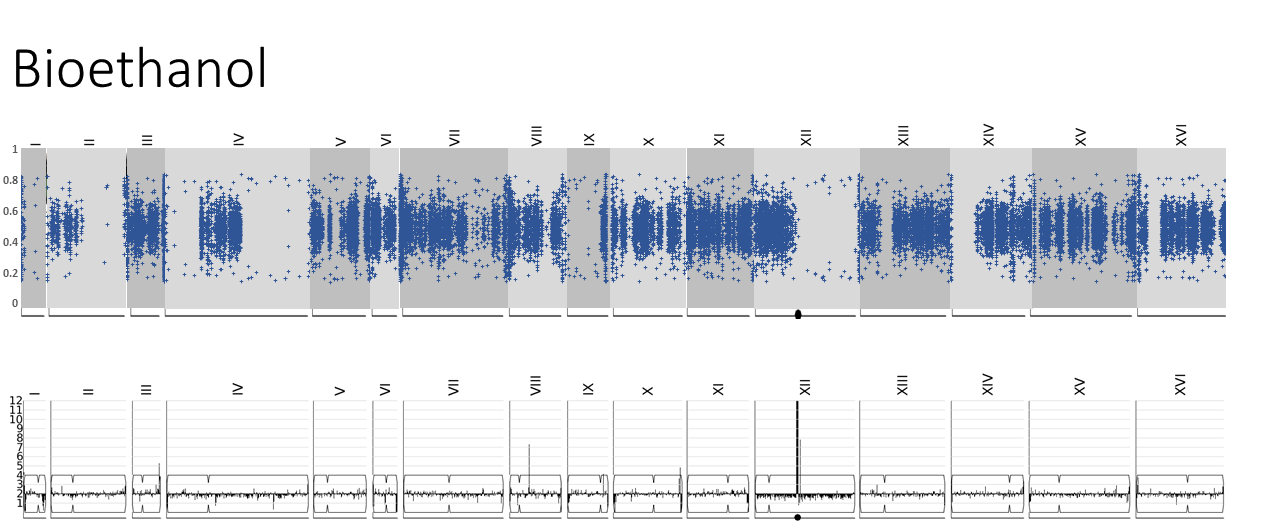

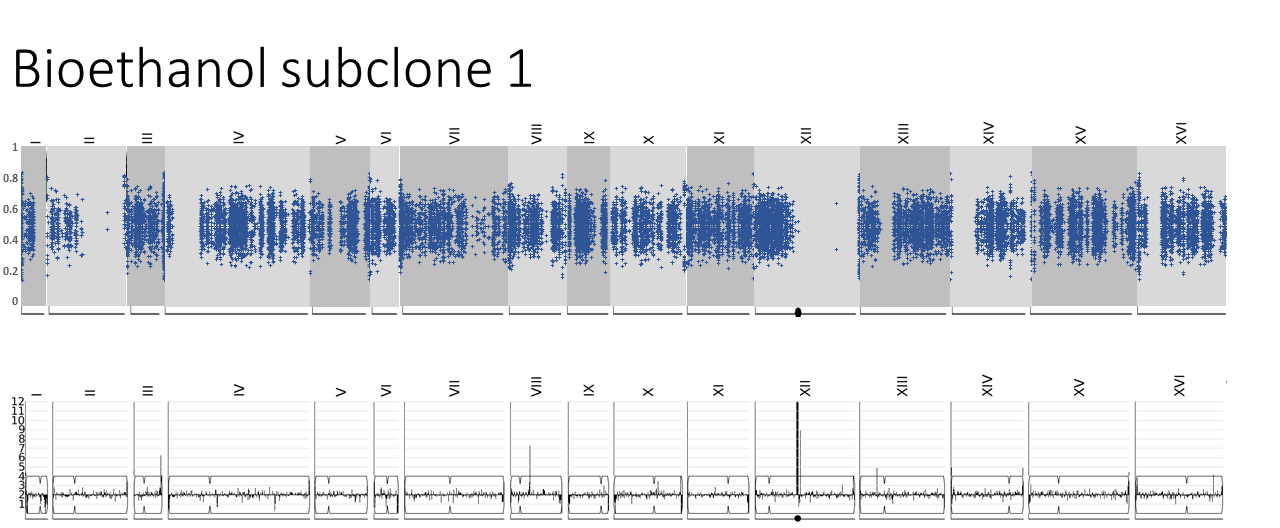

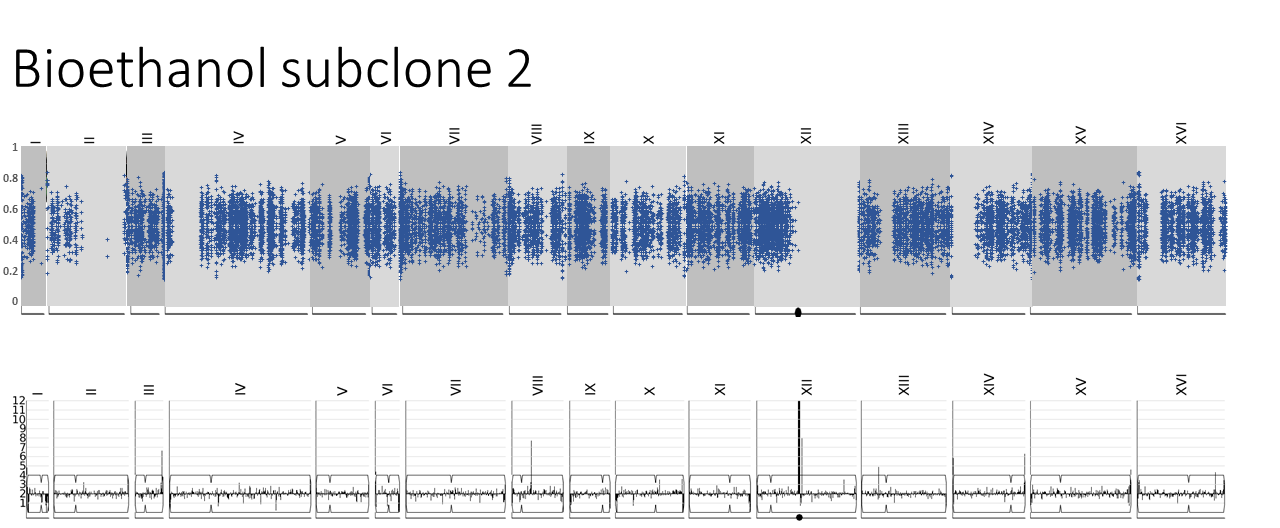

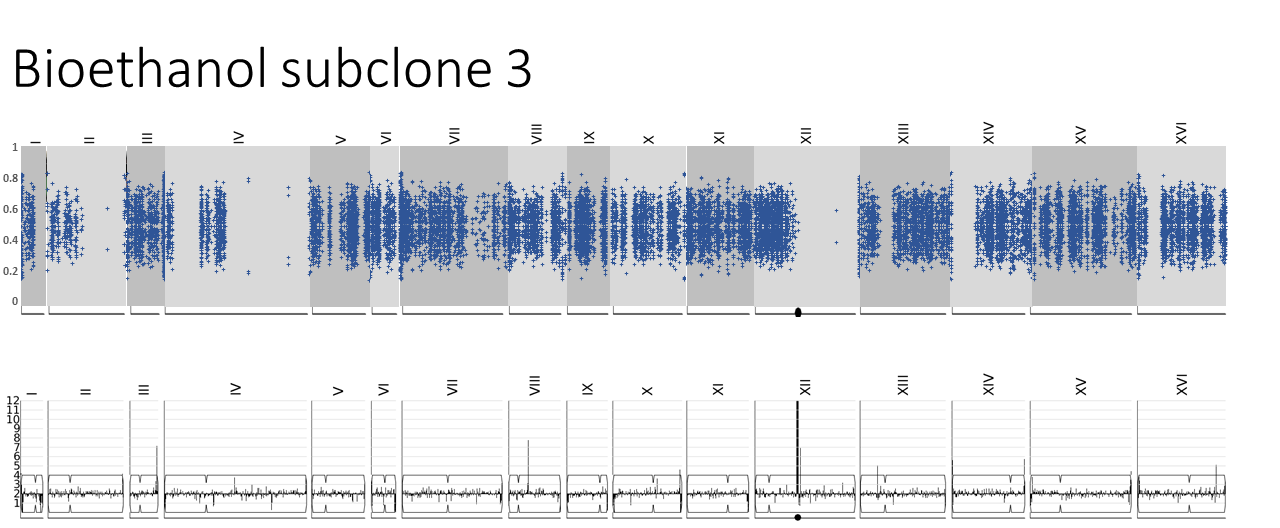

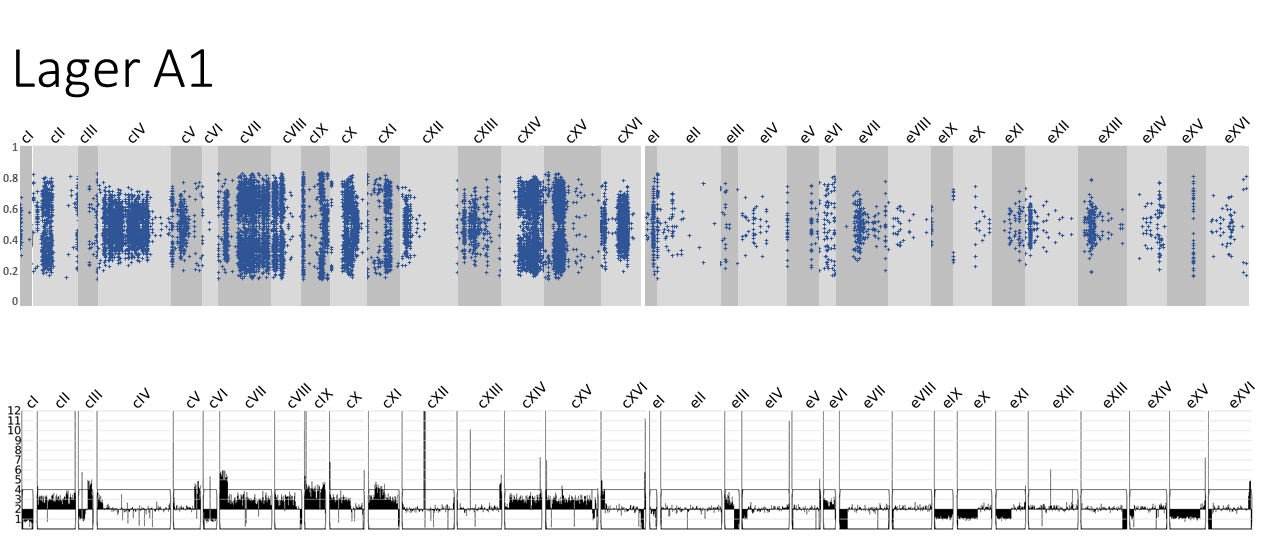

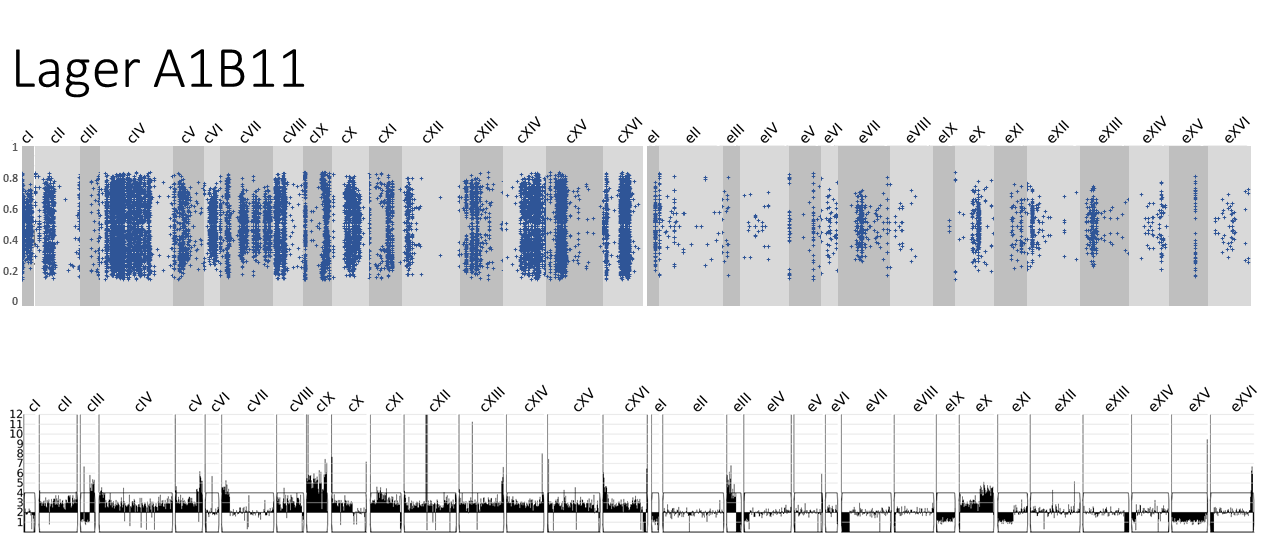

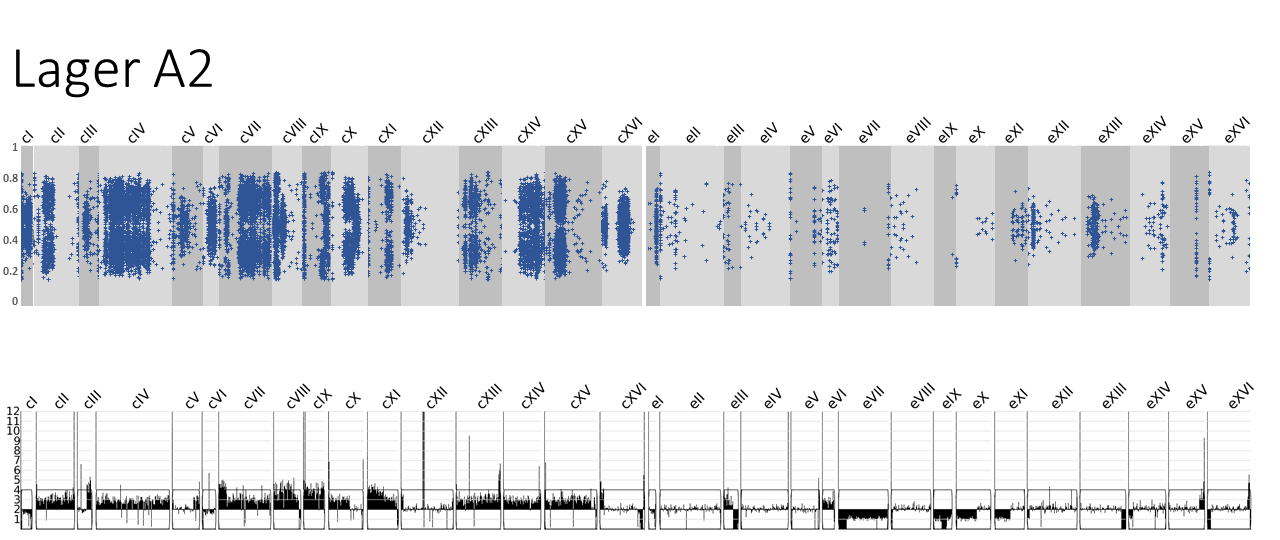

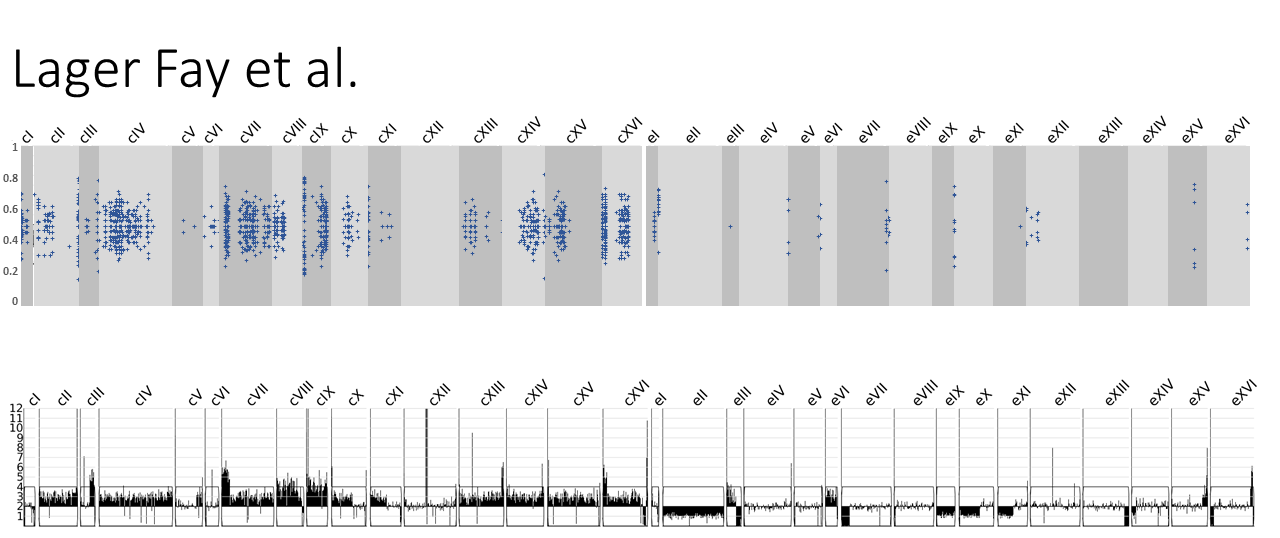

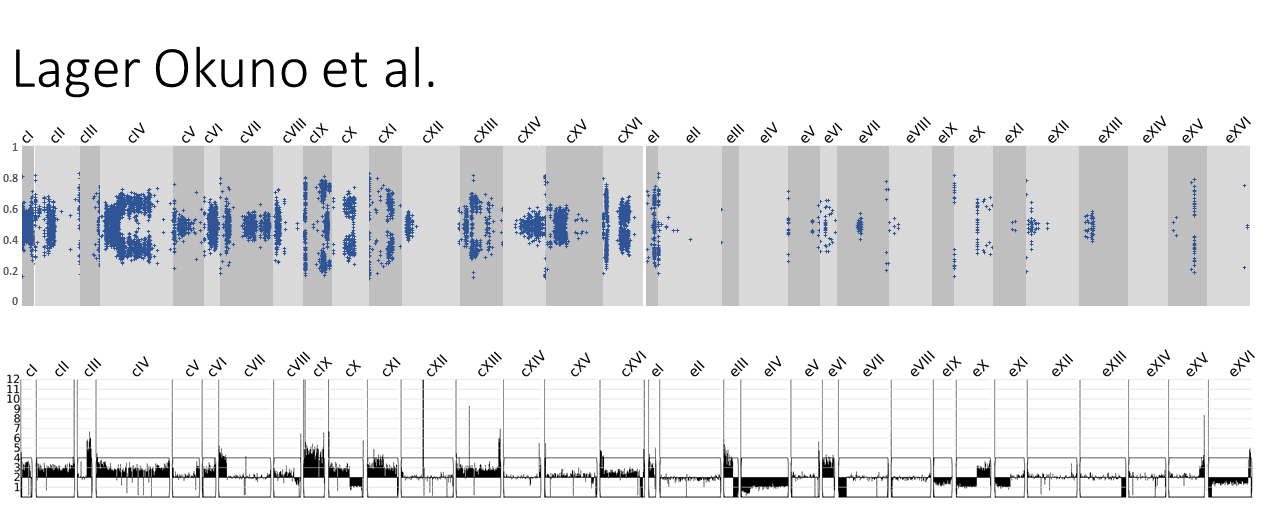

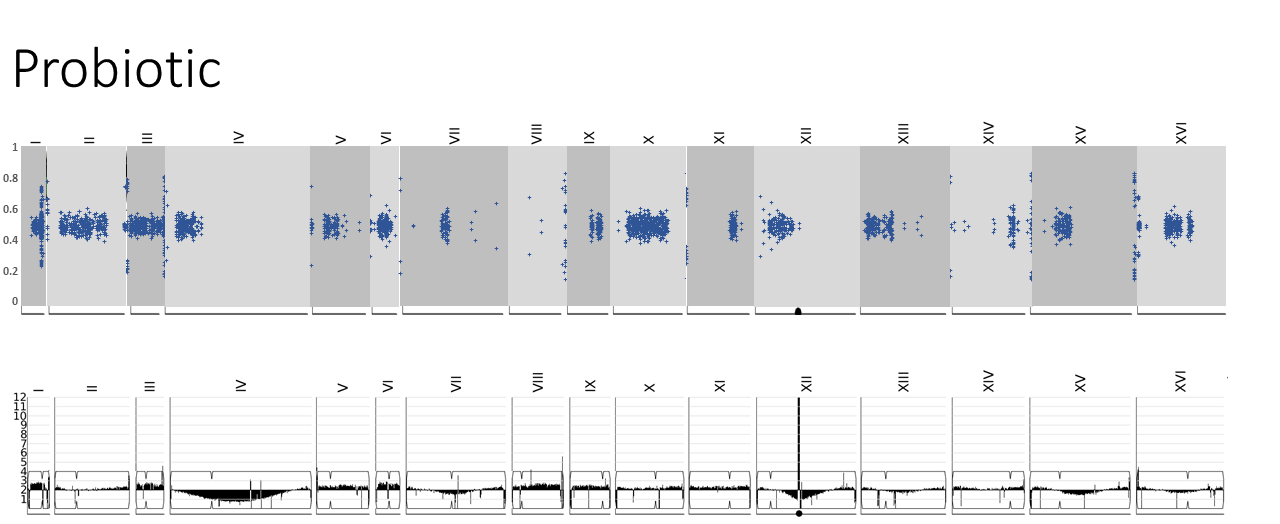

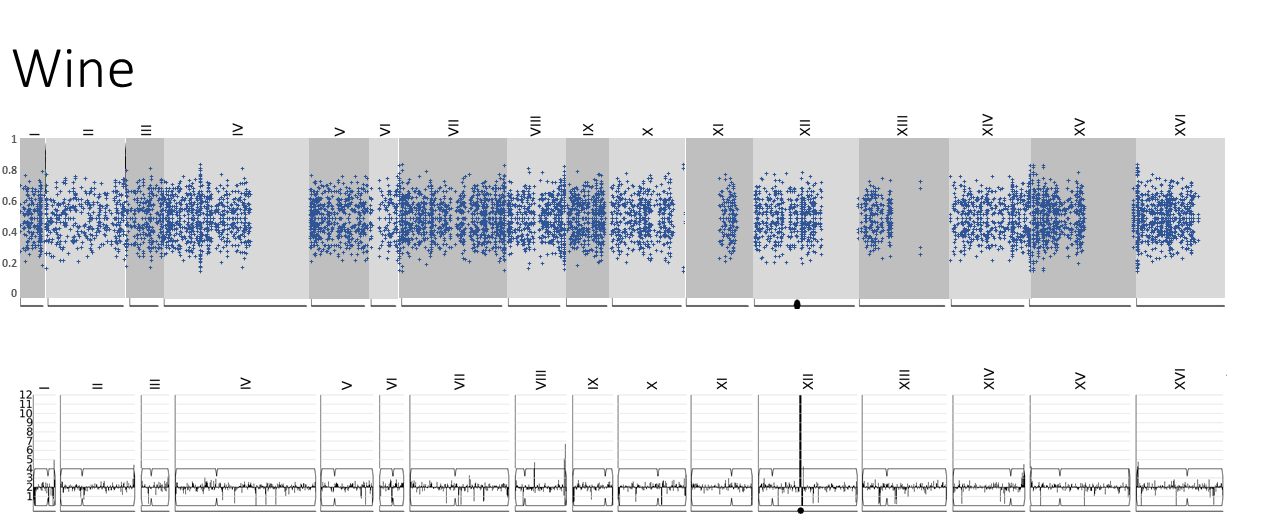

**Figure S2.**

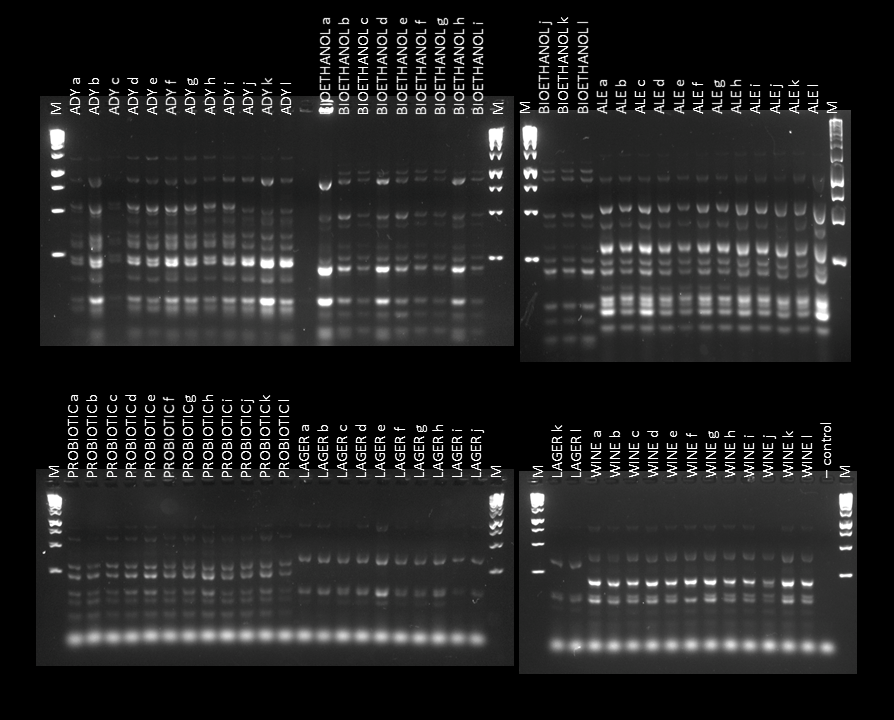

**Figure S3.**

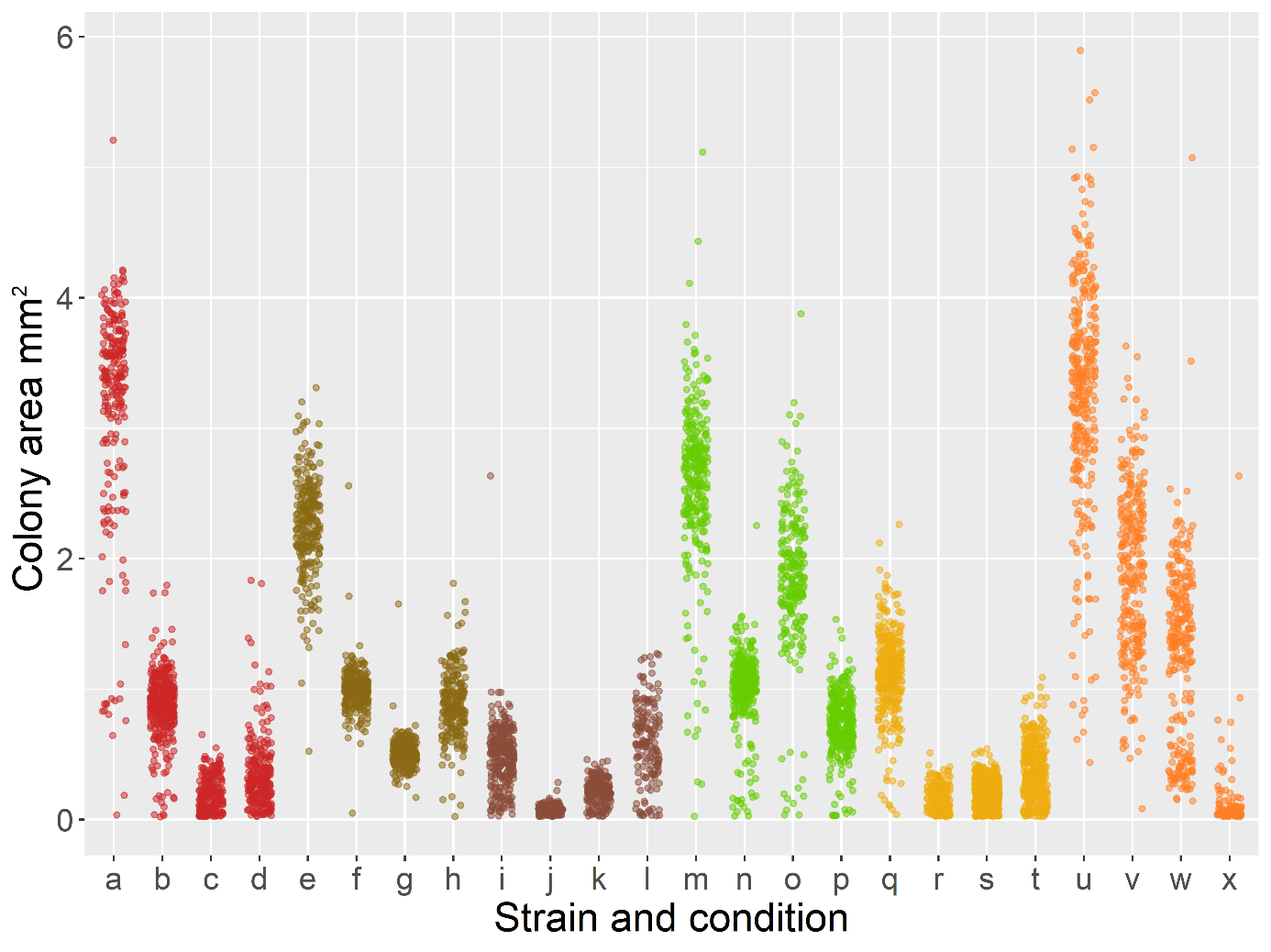

**Figure S4.**

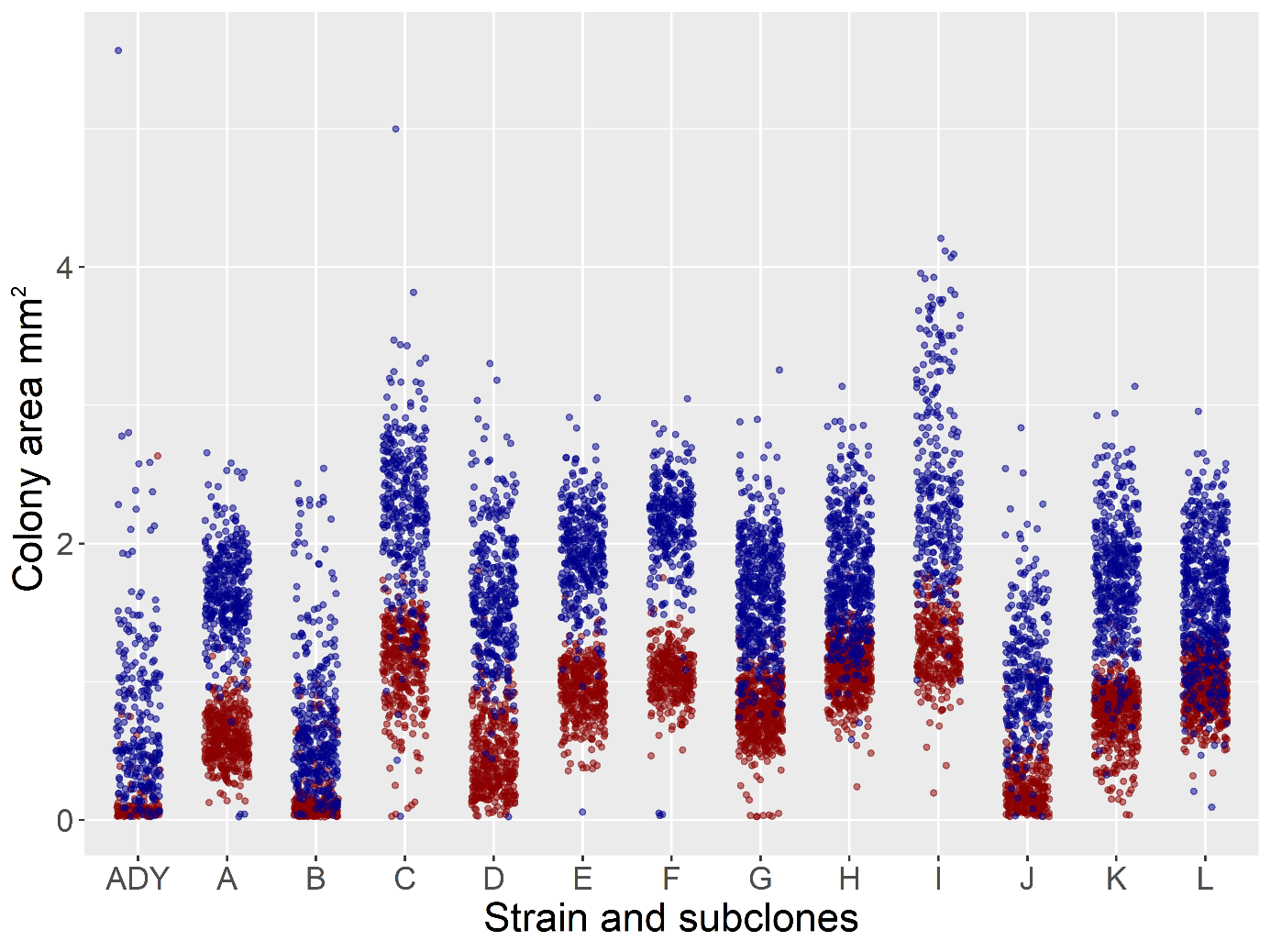

**Figure S5.**

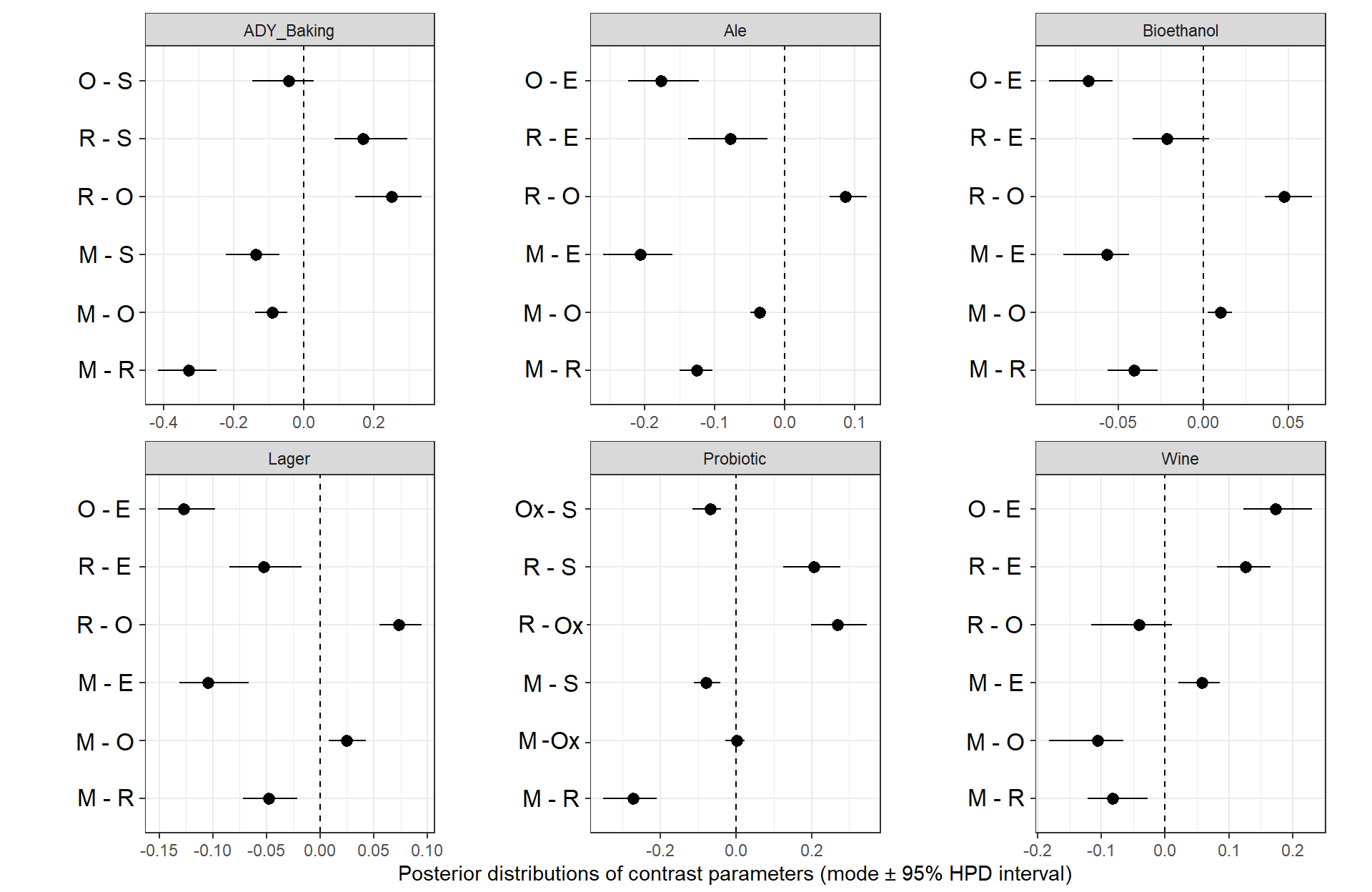

**Figure S6.**

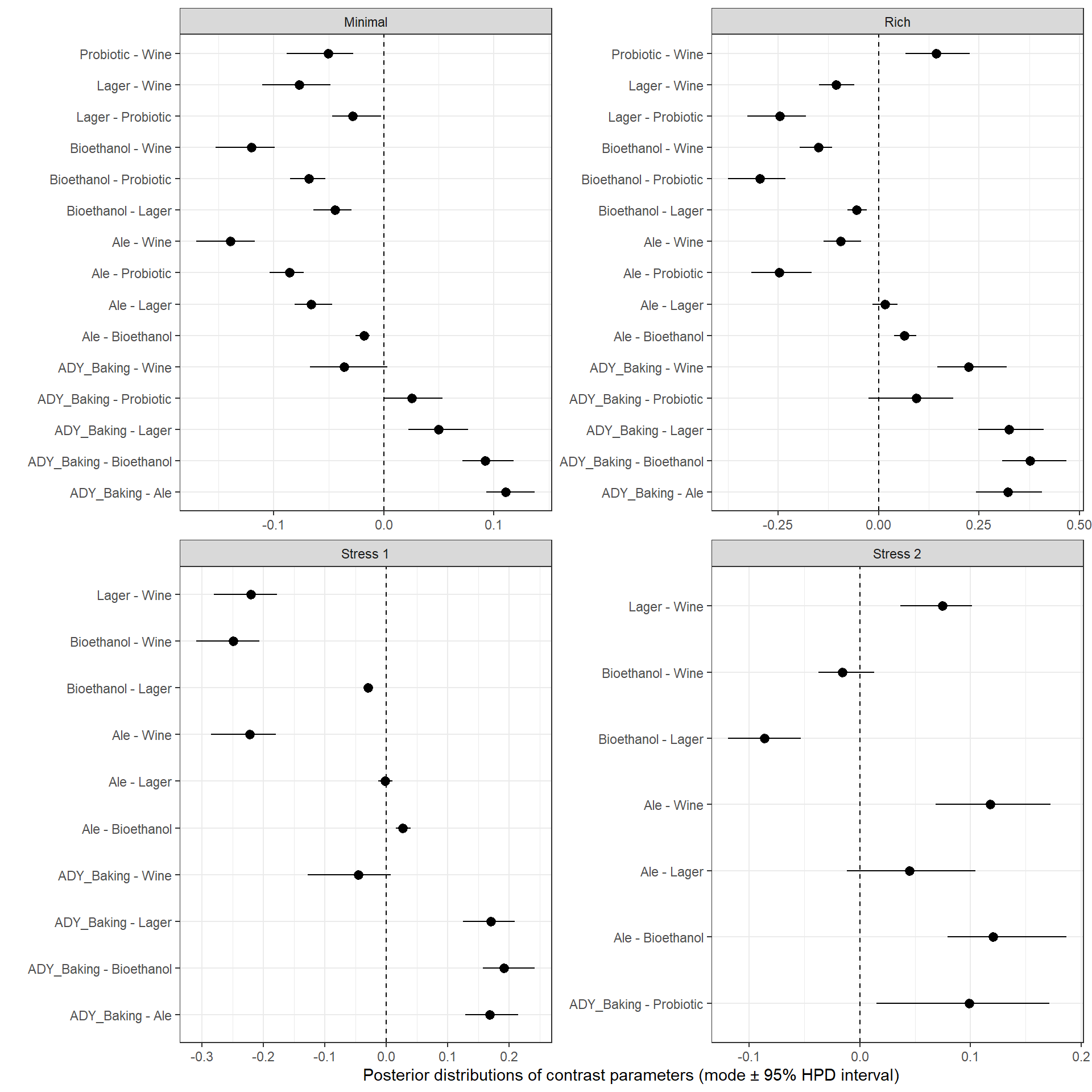

**Figure S7.**

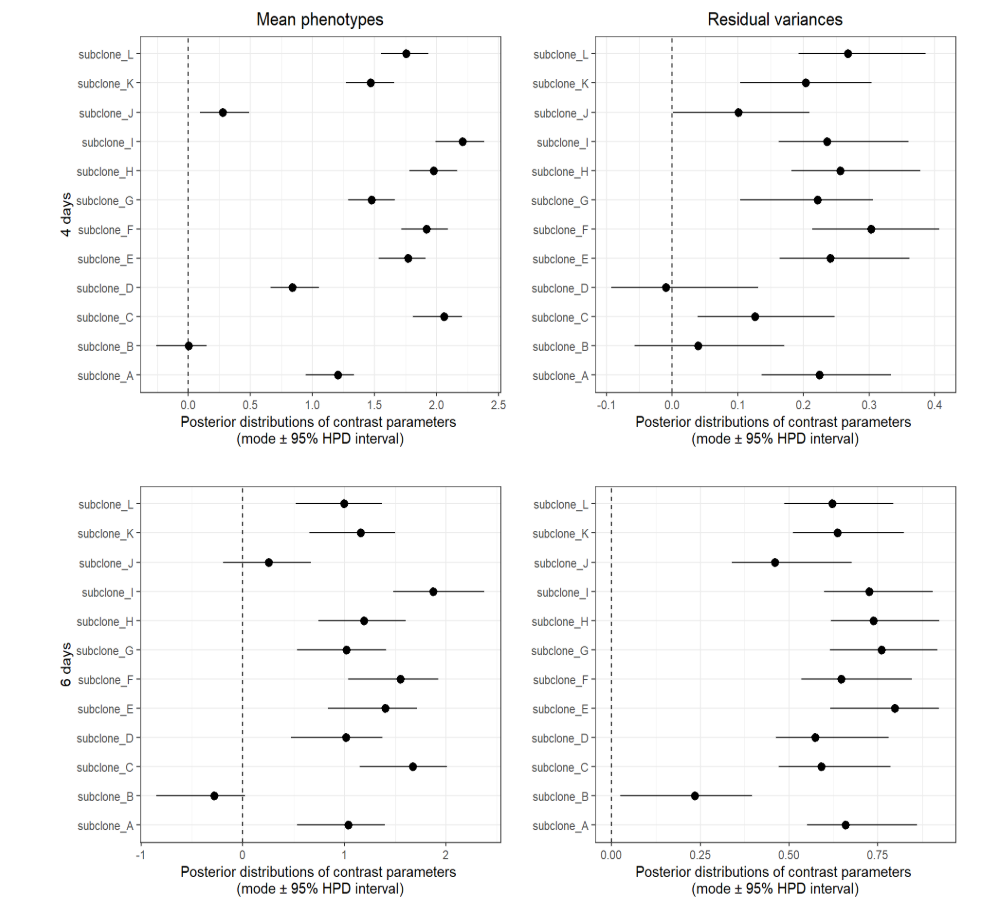
